# Automated eDNA and eRNA Profiling for Biodiversity Monitoring in Marine and Freshwater Ecosystems

**DOI:** 10.1101/2025.10.30.685529

**Authors:** Robert G. Beiko, Jennifer Tolman, Soma Sardar Barawi, Mohamed Fares, Sneha Surya Narayana Murthy, Tom Knox, Connor M. Mackie, Iain Grundke, Nicholas W. Jeffery, Ryan R. E. Stanley, Vincent Sieben, Julie LaRoche

## Abstract

Automated sampling enables the collection and analysis of eDNA from regions that are limited by site access, sampling times, and operator safety. eDNA sampling devices must be rigorously tested against existing technologies to demonstrate fitness across different operational settings and sample quality. The Dartmouth Ocean Technologies, Inc. (DOT) automated eDNA sampler preserves samples and can be deployed at a range of temperatures and depths. The DOT sampler has previously been tested in marine environments for up to three months, with validation against manual protocols.

In this study we tested the DOT sampler in four water bodies in Nova Scotia, Canada, with an expanded set of genetic analyses. We successfully profiled prokaryotes, eukaryotes, and fish using the 16S, 18S, and 12S ribosomal RNA genes respectively, in a brackish pond, a freshwater lake, and two marine harbours. eDNA samples collected by the DOT sampler were statistically concordant with manual Niskin-bottle samples in a range of aqueous habitats. We detected taxonomic groups consistent with the salinity level of each sampled habitat, including invasive species such as smallmouth bass and chain pickerel in the freshwater lake. One marine harbour was sampled at pre-defined time intervals in the days following a significant rainfall event during which site access was limited. We detected ten times as many probable fecal-associated bacteria by proportion at this site relative to the other marine harbour. Onboard preservation of samples in RNAlater allowed the identification of groups with different levels of metabolic activity, and shotgun metagenomic analysis identified key metabolic pathways and a small number of sequences with homology to known antimicrobial-resistance genes.

## Introduction

Effective management of natural ecosystems, whether terrestrial or aquatic, relies on robust information about their status and ongoing changes. Human activities exert complex and often cumulative pressures on ecosystems, impacting the organisms that inhabit them (Halpern et al. 2019). Halting the loss of biodiversity and erosion of ecosystem functioning has become a global priority, as reflected in initiatives like the Kunming–Montreal Global Biodiversity Framework (GBF) and the Biodiversity Beyond National Jurisdiction (BBNJ) treaty. The GBF’s ambitious ‘30 × 30’ target seeks to conserve 30% of land, waters, and seas by 2030. Achieving this goal necessitates approximately a fourfold increase in ocean protection globally (from 8% to 30%) and a doubling of land and inland water conservation (from 16% to 30%) across jurisdictions and governance boundaries (Buenafe et al. 2025; UNEP–WCMC and IUCN 2025). As the global conservation estate expands, so too does the need for effective monitoring tools to track biodiversity change and assess the efficacy of protection mechanisms, such as Marine Protected Areas (MPAs) and Other Effective Conservation Measures (OECMs) (Navarro et al. 2018; Perino et al. 2022). This need is intensified by the accelerating impacts of climate change, which are driving shifts in distribution and ecosystem function within aquatic environments (e.g., Perry et al. 2005; Hochkirch et al. 2021; Hannah et al. 2024; Irvine et al. 2025).

Traditionally, monitoring ecological change in aquatic environments has relied on a wide range of methods tailored to specific environmental and logistical constraints, such as sampling depth, accessibility, and the availability of taxonomic expertise. Manual identification of taxa and biodiversity profiling suffers from several limitations, including the disturbance and collection of organisms *in situ*, and the need to recognize sometimes subtle morphological differences to assign species identities. Furthermore, accurate taxonomic identification often depends on specialized expertise, which has been on the decline (Giangrande 2003) and can itself introduce variability and bias (Turton-Hughes et al. 2024). Traditional collection and identification techniques often lack the ability to generate precise profiles of microbial diversity, are limited in their ability to identify rare and elusive species, and may be infeasible in areas that are remote, unsafe, or where species collection may be unacceptably disruptive (Staley and Konopka 1985; Johnson et al. 2021; Fediajevaite et al. 2021). Environmental DNA (eDNA) sampling has gained considerable attention for its potential to augment or replace traditional methods in aquatic settings (e.g., Deiner et al. (2017); He et al. (2023); Picq et al. (2024)). Shed cells and free DNA can be profiled using a range of techniques to estimate species composition within an environment (Thomsen and Willerslev 2015). This molecular approach overcomes many limitations of visual or physical sampling, enabling comparable protocols across highly divergent ecosystems in a comparatively cost-efficient and non-invasive manner (He et al. 2023). eDNA can identify living and dead organisms depending on shedding rate and DNA stability for a given set of environmental factors (Pochon et al. 2017; Collins et al. 2018, 2021; Chipuriro et al. 2022). Similarly, RNA can be sampled from the aquatic environment. In comparison to eDNA, eRNA is much more unstable in the environment, and emerging analysis techniques provide more of a focus on living organisms and current community structure (Giroux et al. 2022). eRNA can also capture the magnitude of gene expression in different organisms, which can reflect local stressors or responses to environmental change (Yates et al. 2021).

Species identification and biodiversity profiling are often based on analysis of a single gene or gene fragment that is chosen to optimize taxonomic scope and resolution, with the polymerase chain reaction (PCR) used to amplify the desired range of taxa. Highly prevalent genes involved in core cellular functions are frequently used for profiling and identification: for example, small subunit ribosomal RNA genes such as the bacterial 16S gene, the eukaryotic nuclear 18S gene, and the mitochondrial 12S gene can provide a broad taxonomic scope, as can the mitochondrial cytochrome oxidase 1 (COI) gene and the *rbcL* gene in plants (Hebert et al. 2003; Burgess et al. 2011; Thomsen and Willerslev 2015; Stat et al. 2017; Zinger et al. 2019). Non-genic regions such as the intergenic transcribed spacer (ITS) region and the mitochondrial D loop have also been used (Tedersoo et al. 2022). Taxonomic precision can be increased by designing and validating specific PCR primers to target specific groups, at the species or genus level, depending on the aims of the study (MacDonald and Sarre 2017; Zarzyczny et al. 2025). Metagenomic and metatranscriptomic techniques can be used to profile the taxonomic and functional potential of a community by sequencing random fragments of DNA and RNA.

Remote or extreme environments such as the deep sea or offshore regions present significant sampling challenges, often due to limited resources or physical constraints of sampling *in situ* (Levin et al. 2019). Data collection is frequently restricted spatially (e.g., focused surveys within limited areas that can be regularly accessed), temporally (e.g., dictated by suitable weather or seasonal windows), or in sensitive and protected habitats. Autonomous samplers can reduce the need for costly and potentially risky travel to remote locations, for example at high latitudes, altitudes, or the deep ocean (e.g., Cote et al. 2023). Several commercial-grade and research-grade eDNA samplers are available to end users (Yamahara et al. 2025); these differ by degree of autonomy, depth tolerance, prevention of sample cross contamination, filter pore size, and sample preservation. The Robotic Cartridge Sampling Instrument (RoCSI) developed by the National Oceanographic Center (Southampton, UK) is fully autonomous with a belt-loading configuration of filters. The Ocean Diagnostics Inc. sampler, Ascension, is suitable for semi-autonomous operations and achieves high flow rates by using larger 5-micron pore size filters, but does not preserve samples *in situ*. The 45 kg Smith-Root autosampler (SR-eDNA) has been deployed alongside rivers and lakes and can collect up to eight samples on a schedule using self-preserving filter packs for rivers and lakes (George et al. 2024). The Dartmouth Ocean Technologies Inc. (DOT) eDNA sampler (Hendricks et al. 2023) has a capacity for nine filters with different pore-size options, is fully autonomous with preservation capabilities, and uniquely uses acid cleaning between samples to prevent cross-contamination. Prior to large-scale deployment and use, autonomous samplers must be rigorously tested to ensure they can produce results similar to those generated using conventional eDNA sampling approaches involving manual filtration. They must also be capable of preserving samples during long-term deployments with minimal risk of sample decay, contamination of components, biofouling, and mechanical failure.

The DOT sampler has previously been compared to other methods of water sampling, such as Niskin bottles, in multiple field studies (Hendricks et al. 2023). During a nine-week test deployment in Halifax Harbour, the programmed DOT samplers performed similarly at detecting marine fish diversity relative to water samples collected each week and filtered on a standard vacuum pump (Van Wyngaarden et al. 2024). However, the invertebrate communities detected using the COI marker differed between methods, which were attributed to differences in filter pore size (0.22μm vs 1.2μm for the DOT sampler and freshly collected samples, respectively). Notably, DNA concentrations preserved *in situ* by the DOT sampler showed no evidence of DNA degradation over 9 weeks, which was comparable to preserving the fresh sample filters at –80°C. Biofouling from algae and encrusting animals was also minimal over this time period, despite the sampler’s placement in a shallow, high-traffic industrial port environment where biofouling is typically prevalent. These results suggest that long-term use of the DOT sampler is feasible in coastal marine environments.

While autonomous eDNA technologies such as the DOT eDNA samplers represent a significant advancement for aquatic biodiversity monitoring, its performance has not yet been comprehensively assessed across the full range of environmental contexts in which it is needed, such as freshwater, estuarine, and marine systems, or sites with varying levels of disturbance, salinity, or biofouling risk. Moreover, although studies have explored the concurrent use of eDNA and eRNA for assessing community diversity (e.g., Laroche et al. (2017); Giroux et al. (2022)), few have systematically evaluated the ability of autonomous samplers to preserve and recover both nucleic acid types from the same samples.

We deployed a DOT sampler in multiple aquatic settings and performed comprehensive biodiversity profiling from preserved eDNA and eRNA. While the primary objective of our study was to test the DOT sampler’s ability to collect and preserve a combination of eDNA, eRNA, and microbial DNA for metagenomics, we also aimed to identify prokaryotes and eukaryotes related to human and ecological health. Deployments were carried out in a brackish pond, a freshwater lake, and two marine harbours in Nova Scotia, Canada. Biodiversity profiles inferred from samples collected using the DOT sampler were comparable to those obtained from Niskin bottles, with the exception of ascomycete fungi in the brackish pond. Parallel analysis of eDNA and eRNA revealed distinct community assemblages, consistent with temporal variation in biodiversity. We successfully identified native and invasive species and demonstrated the ability to detect shifts in microbial diversity after a disruptive rainfall event in Halifax Harbour. Our results demonstrate the effectiveness of autonomous sampling in multiple habitats and the potential for deployments in settings where frequent manual sampling may be impractical for reasons of safety and cost.

## Methods

### The DOT eDNA sampler

The DOT eDNA sampler is an autonomous sampling system capable of long-term deployments in remote, difficult-to-reach locations as described in (Hendricks et al. 2023). This sampler consists of 3 subsections: a filter cassette that holds 9 discrete 25 mm filters with 0.22 μm polycarbonate (PC) membrane, an instrument housing, and a reagent housing. The filter-cassette design allows rapid removal and replacement of filters in the field by operators. The second subsection, the instrument housing, contains the 26 valves, pump, pressure sensors, control circuitry, and inner fluid lines required for automating sample cleaning, collection, and preservation. The final subsection is the reagent housing, which is a free-flooded chamber that contains three separate IV bags that are preloaded with a) RNase– and DNase-free water, b) 5% HCl, and c) RNAlater preservative. A fourth bag is initially empty and is used to capture waste reagents during the acquisition and preservation of the samples. The RNase– and DNase-free water is used for priming the system lines and filters when new filter cartridges are installed. The 5% HCl is used to sterilize and decontaminate the sample inlet and inner tubing before a sampling event. The RNAlater preservative is loaded into filters after the sampling event to preserve both RNA and DNA. The waste bag is used to capture the preservative to ensure it does not enter the environment.

### Sample Collection

Water samples were collected at four locations in Nova Scotia (Figure 1) in July 2023: Sheeps Head Pond (SHP: Tantallon, NS) on July 17, a jetty in Lunenburg Harbour (Lun) on July 19, Grand Lake (GL: Oakfield, NS) on July 21, and a jetty at the Centre for Ocean Venture and Entrepreneurship in Halifax Harbour (CJ: Dartmouth, NS) on July 22-24 (Table 1, Table S1). Sampler deployment was carried out in a similar manner to Hendricks et al. (2023), with all handling of filters done using sterile technique with gloves. Prior to deployment, field equipment was cleaned and DNA filters loaded into the filter cassette holder, which connects to the fluid manifold and syringe pump in the sampler. Samples were collected from SHP, Lun, and GL at approximately 45-minute intervals between 9:30 AM and 4 PM, with minor variation between sites. Sampling at CJ consisted of nine automated sample collection and preservation runs performed at four-hour intervals between 5:27 PM on July 22 and 12:02 AM on July 24. After retrieval, preserved filters were physically removed from the cassette, LUER caps were installed on the inlet/outlet ports of each filter, then samples were bagged, taken to the laboratory, and stored frozen at –20C until further processing. The sampler was powered by a DOT battery pack in all four locations, with a communication cable run to a laptop on shore for monitoring. The sampler was deployed by rope in three of the four locations: in SHP, to a depth of 1 metre from the back of a small inflatable vessel; in GL, to a depth of 1.5 metres from a private dock; and in Lun, to a depth of 1.5 metres from a commercial dock. In SHP, GL, and Lun, a Niskin bottle was lowered by rope into position next to the eDNA sampler and filled with sample water from the same location. A manual eDNA sample was then collected via a small peristaltic pump into an additional filter, with preservation post filtering. For the CJ samples, the DOT sampler was deployed via rope to a depth of 1.5 metres using drop boards attached to the dock near the sewage outflow pipe.

**Figure 1.**
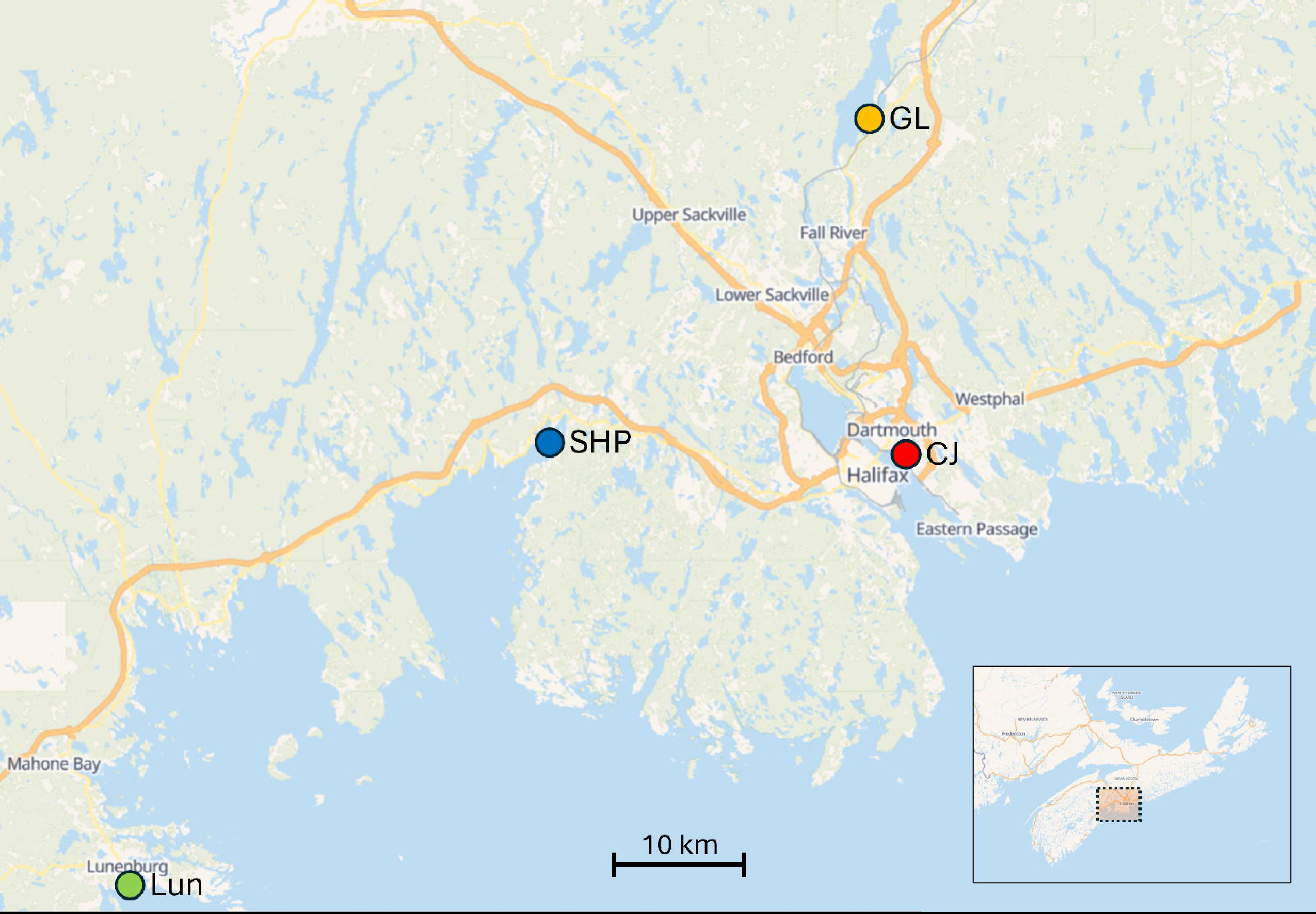
The four sampling locations in this study. CJ = Cove Jetty, GL = Grand Lake, Lun = Lunenburg, SHP = Sheeps Head Pond.

**Table 1.** Summary of samples collected at each site, sampler, and sequencing marker. Int = interval in minutes or hours, Vol = sampled volume in mL. DOT and Niskin samples were relatively low in volume filtered, likely due to the high particulate content of the samples, indicating manual and automatic methods rapidly loaded the filter membrane.

| Site | Sampler | Number of samples |  |  |  | Int | Vol |
| --- | --- | --- | --- | --- | --- | --- | --- |
|  |  | 16S | 18S | 12S | MG |  |  |
| SHP | DOT | 7 | 7 | 2 | 0 | 45m | 39.4 |
| SHP | Niskin | 7 | 7 | 2 | 0 | 45m | 42 |
| Lun | DOT | 9 | 9 | 2 | 0 | 45m | 99.1 |
| Lun | Niskin | 9 | 9 | 2 | 0 | 45m | 28 |
| GL | DOT | 9 | 9 | 2 | 0 | 45m | 40.7 |
| GL | Niskin | 9 | 9 | 2 | 0 | 45m | 52 |
| CJ | DOT | 9 | 9 | 2 | 9 | 4h | 108.1 |

### DNA / RNA Extraction and Sequencing

eDNA was extracted from frozen filters using the DNeasy Plant Mini Kit (Qiagen, Germany) with a modified lysis procedure, including a five-minute incubation at room temperature with 50µL of 5mg/mL lysozyme (MP Biomedicals, USA) followed by 1h at 52°C in 400µL Buffer AP1 supplemented with 45µL of 20mg/mL proteinase K (Fisher BioReagents, United Kingdom). Extraction then proceeded following the manufacturer’s instructions, and eDNA was eluted in 50µL of Buffer AE. Two filters from each sample set were processed for eDNA and eRNA using the AllPrep® Micro Kit (Qiagen, Germany). Cells were lysed via a 5min incubation at room temperature with 50µL of 5mg/mL lysozyme (MP Biomedicals, USA) followed by 15min at 52°C in 600µL Buffer RLT+ supplemented with 45µL of 20mg/mL proteinase K (Fisher BioReagents, United Kingdom) and 12µL of 2M dithiothreitol (Invitrogen, USA). An on-column DNase digestion was performed using the RNase-free DNase set (Qiagen, Germany). Cleaned, purified eRNA was eluted in 20µL of RNase-free water; eDNA was eluted in 50µL of Buffer EB. Nucleic acid concentrations and purity were assessed via a NanoDrop2000c (Thermo Scientific, USA). 6µL eRNA were treated with ezDNase and reverse-transcribed to cDNA using Superscript IV VILO (Invitrogen, USA) according to the manufacturer’s instructions; no-RT reactions were performed alongside to confirm the absence of any contaminating eDNA.

An amplicon sequencing strategy was designed to cover ecosystem biodiversity through detection of prokaryotic and eukaryotic sequences. For all eDNA and cDNA samples (RT and no-RT), the metabarcoding assays targeted regions of the prokaryotic and eukaryotic rRNA genes, 16S (V4-V5 regions; Parada et al. 2016), and 18S (V4; Comeau et al. 2017), respectively. For a subset of samples, specific regions of eukaryotic mitochondrial (mt) genomes were targeted for fish (12S miFish; (Miya et al. 2015; Stoeckle et al. 2022)) specifically. eDNA and cDNA for amplicon sequencing of 16S rRNA V4-V5 and 18S rRNA V4 regions were sent directly to the Integrated Microbiome Resource (IMR) facility at Dalhousie University and processed as described by (Comeau et al. 2017). PCR reactions targeting the 12S gene were performed in 20µL reactions using HotStar Taq (Qiagen, Germany), 250nM forward and reverse primers (IDT, USA), 200µM dNTPs (Invitrogen, USA), and 2µL eDNA template. 12S reactions used 4µg bovine serum albumin (NEB, USA) with touchdown cycling parameters of 15 min at 95°C, 13 cycles of [30s 94°C, 30s 69.5°C-1.5°C/cycle, 90s 72°C], 30 cycles of [30s 94°C, 30s 50°C, 45s 72°C], and 10 min at 72°C. PCR products were purified with the Monarch spin PCR & DNA cleanup kit (NEB, USA) and submitted to the IMR for Illumina MiSeq sequencing (2×300bp paired end).

Shotgun metagenomic sequencing was performed by the IMR for all 9 CJ samples. Libraries were prepared using the Illumina Nextera Flex kit as in (Comeau and Filloramo 2023) and sequenced at 2X depth on the Illumina NextSeq2000 (2×150bp paired end).

### Data Analysis

All amplicons derived from environmental DNA and RNA were processed the QIIME2 software package v2024.5.0 (Bolyen et al. 2019). All amplicons were trimmed and denoised using DADA2 (Callahan et al. 2016). Taxonomic classification of 16S and 18S sequences was performed using the Naïve Bayes classifier with v138.1 of the SILVA database (Quast et al. 2013). Only taxonomic assignments with a confidence score greater than 0.7 at a given rank were kept. For 12S, taxonomic assignment was performed using BLASTN against the MIDORI2 reference databases vGB266 (Leray et al. 2022) with all unique haplotype sequences included.

In the initial analysis, samples were rarefied to 1000 sequences each in order to maximize the number of retained samples. The Shannon diversity of all samples was computed and compared using a paired-sample *t*-test and a significance threshold of 0.05. Aitchison distances were calculated between all samples and used as the basis for principal coordinates analysis, both with the full dataset and with each site considered individually. Taxonomy-based analyses (barplots, UpSet plots, heatmaps, and time-series visualizations) were performed using an abundance threshold *T*. In any given analysis, a taxonomic group at the appropriate rank with abundance ≥ *T* in at least one sample was included. The choice of rank and *T* was analysis-specific.

Each sampled location had two time points where both 16S and 18S DNA and RNA were sequenced. Although the small sample size precluded statistical analysis, we compared DNA and RNA samples in a way that accounted for the compositional nature of amplicon sequence data. We first computed the centered log ratio (CLR) for each taxon *t* in each sample *s* using the standard formula:

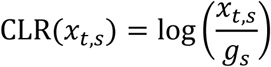

where *x*_*t*,*s*_ is the relative abundance of taxon *t* in sample *s*, and *g*_*s*_ is the geometric mean of the relative abundance of all taxa in *s*.

We then computed the CLR difference *D*_*t*,ℓ_ for each taxonomic group *t* and location ℓ as follows:

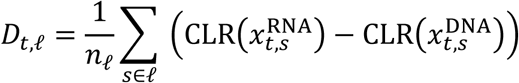

In our case, *n*_ℓ_ = 2 (two paired DNA–RNA samples per location). We performed this comparison only on taxa that had nonzero abundance across all four samples at a given location, and further required that a taxon be present with a minimum abundance of at least *T* in one of the four samples.

For the metagenomic data generated from the 9 CJ samples, quality assessment and statistical analysis of raw reads was performed using FastQC v.0.12.1 (Andrews 2010). Subsequently, reads were processed using FastP v0.24.0 (Chen et al. 2018) to remove adapters, duplicates, and trim low-quality bases, with reads shorter than 50 bp after trimming discarded. The resulting high-quality reads were used for all subsequent downstream analyses. Assemblies were generated separately for each of the nine samples using MEGAHIT v1.2.9 (Li et al. 2015) with the ‘–meta-large’ preset (k-mer sizes 27–127 bp, step size 10). Assembled contigs shorter than 1000 bp were discarded. Assembly quality was evaluated using QUAST v5.2.0 (Gurevich et al. 2013) with the MetaQUAST extension, generating metrics including the number of contigs, total assembly length, largest contig, and N50.

Taxonomic classification of processed reads and contigs was performed with Kraken2 v2.1.2 using default parameters (Wood et al. 2019). Functional profiling was carried out with HUMAnN v3.8 (Beghini et al. 2021), which assigned annotations for MetaCyc reactions, Gene Ontology (GO) terms, and KEGG Orthology (KO) identifiers. These annotations were further processed using KEGG-decoder v1.3.0 (Graham 2020) to assess the presence and completeness of metabolic pathways. Detection of antimicrobial-resistance genes (ARGs) was performed using the Resistance Gene Identifier (RGI) v6.0.5 on contigs (Alcock et al. 2023). RGI Perfect and Strict matches were then compared against the NCBI clustered non-redundant protein database (2025/07/05 release) using the BLASTP version 2.16.0 through the NCBI online portal (National Center for Biotechnology Information 2025) with default parameters.

## Results

All DOT and Niskin samples were collected in the prespecified time intervals, although the recovered volumes in some cases were less than 30 mL (Table 1, Table S1). A major disparity was seen in the average volumes obtained by the DOT sampler from the marine sites (Lun = 99.1 mL, CJ = 108.1 mL), versus the brackish and lake sites (SHP = 39.4 mL, GL = 28 mL), likely due to higher particulate loads in SHP and GL rapidly loading the filter membrane in both samples. Despite these small volumes, DNA and cDNA sequence data were obtained from all samples apart from two Niskin samples from Sheeps Head Pond, which generated a single sequence each. Two DOT samples generated fewer than 200 sequences (179 18S sequences from one SHP time point, and 188 16S sequences from one GL time point); these samples were retained in the analysis for comparative purposes. Sequencing of 16S, 18S, and 12S DNA from all DOT samples yielded an average of 24,634 amplicons per sample (n = 76), versus 18,650 for the Niskin samples (n = 50). AllPrep® extractions yielded an average of 94,931 raw sequences per sample obtained from 16S and 18S cDNA sequencing of DOT samples, versus 74,984 for the Niskin samples. Metagenome sequencing of DOT COVE Jetty samples yielded an average of 16,680,966 reads across nine samples.

### Comparisons by Location, Sampler, and Sequence Type

Principal coordinates analysis of DOT and Niskin samples based on Aitchison distance showed very clear distinctions among sites using both 16S and 18S markers, with the first two principal coordinates explaining 53% of total variance for 16S and 41.2% of total variance for 18S (Figure 2). The two marine sites, Lun and CJ, were similar to one another in the 16S samples, while in the 18S samples the two sites were not separated on the first coordinate but were on the second. We then compared sampler type (DOT vs. Niskin) and sequence type (eDNA vs. eRNA) for each site. Sample location, sample type, and sequence type were split along the first two principal coordinates for each of 16S and 18S (Figure S1) markers, although the DOT and Niskin samples were not separated along either of the two principal coordinate axes for the 16S freshwater and brackish samples (SHP and GL). In both cases, the first axis separated eDNA from eRNA samples. The first axis split by sampler for the 16S Lun and all three 18S locations where both sample types were collected (SHP, GL, Lun). The split between DOT and Niskin samples was especially strong in the 16S and 18S Lun samples (PC1 variance explained = 36.3% and 43.3% respectively), and in the SHP 18S samples (PC1 variance explained = 43.3%). The dispersion was larger for DOT than for Niskin samples for 16S and 18S profiles of Lun. Patterns in ASV Shannon diversity differed by site (Figure S2), with the DOT sampler showing higher diversity than Niskin in Lun (16S: p = 0.0005; 18S: p = 0.0009), but lower in GL (16S: p = 0.036; 18S: p = 0.001) and no difference in SHP (16S: p = 0.876; 18S: p = 0.806).

**Figure 2.**
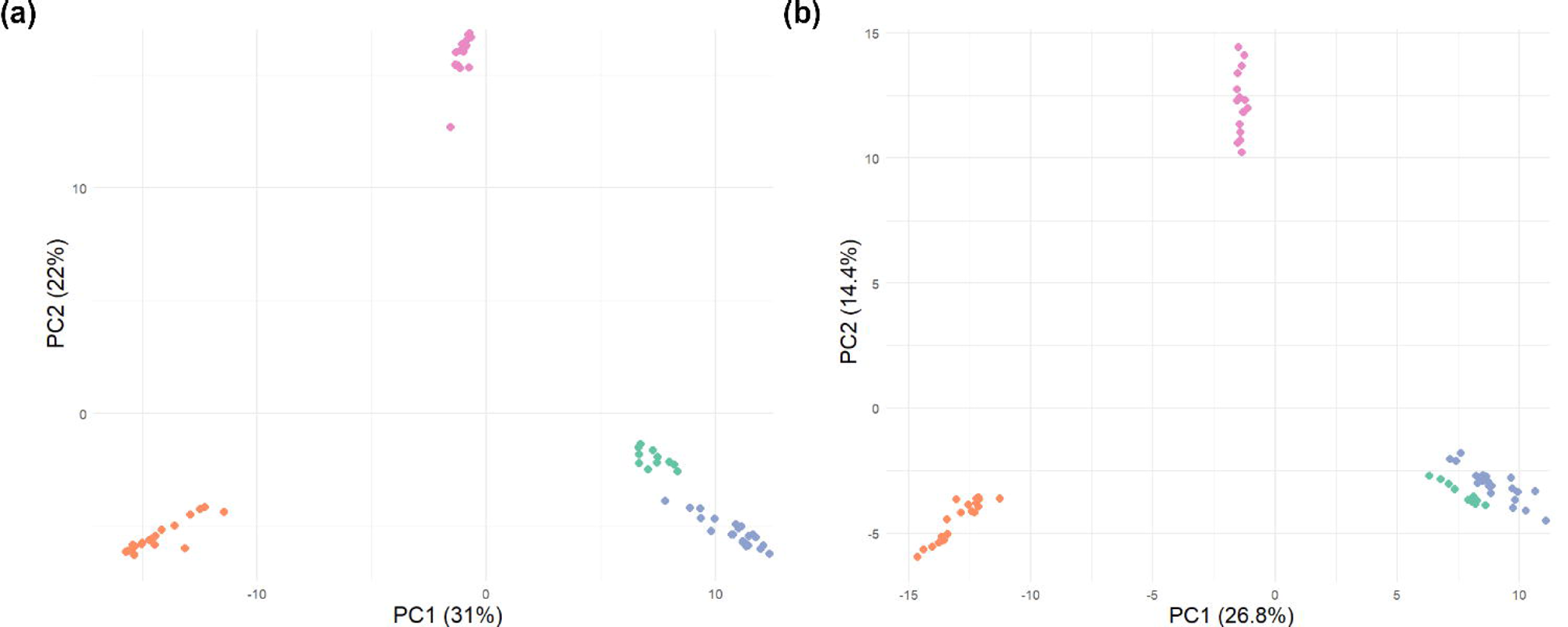
Principal Coordinates Analysis of DOT and Niskin samples. Ordinations of 16S (a) and 18S (b) samples are shown. Location colours are consistent with Figure 1.

Hierarchical clustering of samples based on profiles of common taxa tended to group sites from the same location (Figure S3). Samples from all four sites were distinct in the 16S and 12S samples, while the two marine sites and SHP/GL were intermingled in the 18S profiles. Consistent with the ordination plots, samples from the two marine sites were most closely linked with each other. DOT and Niskin samples for each site were interleaved in the 16S profiles, whereas the DOT marine samples clustered together, with some interleaving of DOT and Niskin samples between SHP and GL. DNA and RNA profiles were generally interleaved for both markers. No taxon was found with >5% abundance at all four sites, with the bacterial families Comamonadaceae, Saprospiraceae, and Alteromonadaceae, several eukaryotic phyla including Ciliophora, Dinoflagellata, and Cryptophyceae, the American eel *Anguilla rostrata* (12S: SHP, GL, Lun), and the alewife *Alosa pseudoharengus* (12S: GL, CJ, Lun) found above this threshold at three sites. Many prokaryotic families and eukaryotic phyla were present across all four sites at a minimum abundance of 0.1%. Other than these, several taxa were present at a minimum threshold of 5% in both CJ and Lun, including the rock gunnel *Pholis gunnellus* and winter flounder *Pseudopleuronectes americanus*.

### Sheeps Head Pond

Sheeps Head Pond is located on a small island in Tantallon, Nova Scotia situated in the headwaters of St. Margarets Bay. Within the island is a small body of water that was originally an inlet of the ocean but was cut off by the construction of causeways in the mid-20th century. Although the pond remains connected to the ocean via culverts, the pond freezes during the middle of the winter. Local residents have expressed concerns about the safety of the pond and sulfurous smells in spring and fall, suggesting the potential presence of sulfate-reducing bacteria.

Observed taxa tended to be found in most or all DOT DNA and RNA samples: 25 families from 16S and 10 species from 12S were always present (Figure S4). The exception was the phylum distribution from 18S, where nine phyla were observed in all DNA and RNA samples except sample SHP-1, in which only 179 18S sequences passed quality-control filters. However, many taxonomic families inferred from 16S markers were limited to a single sample, with samples 6, 2, and 7 containing 11, 8, and 6 unique families respectively. When considering taxa with a minimum abundance of 5% in at least one sample, no 16S families were restricted to a single DOT or Niskin sample (Figure S5). The four RNA samples (2 x DOT, 2 x Niskin) were most similar to each other, with underrepresentation of families including Rhizobiaceae and Comamonadaceae relative to the DNA samples. DOT and Niskin samples were not differentiated in the 16S profiles.

Conversely, there was complete separation between DOT and Niskin samples in the 18S phylum data, with DOT time point 1 as an outlier. Ascomycota were overrepresented in the DOT samples, while animal taxa including Arthropoda, Rotifera, and Platyhelminthes were underrepresented relative to the Niskin samples. We compared the abundance of RNA with DNA for each phylum in the 16S and 18S samples, applying the centered log-ratio (CLR) transformation to correct for the compositionality of the data. In the 16S profiles, phylum Nanoarchaeota was underrepresented in the RNA samples, while Cyanobacteria, Proteobacteria, and Bacteroidota were overrepresented (Figure S6). In the paired 18S samples, Cnidaria were underrepresented, while Cercozoa, Prymnesiophyceae, Diatomea, and SA1-3C06 were overrepresented.

The top 16S families by relative abundance across DOT samples were Cyanobiaceae (19% average abundance), Alcaligenaceae (15%), and Rhodobacteraceae (13.4%) (Figure 3a). To assess the presence of potential sulfate-reducing bacteria (SRBs), we examined the distribution of families specifically within the recently constituted phylum Desulfobacterota (Waite et al. 2020), which includes many SRBs. This phylum accounted for an average of 0.297% of all sequences: several families and orders were observed across time points, with Desulfocapsaceae and Bradymonadales comprising 75.7% of all sequences assigned to this phylum (Figure S7a). While most of the identified families contain sulfate-reducing bacteria, Bradymonadales is not directly associated with hydrogen sulfide production (Wang et al. 2015). Consistent with the heatmaps, the distribution of phyla inferred from 18S sequences was dominated by Ascomycota (average abundance = 63.2%), especially in the first time point. The classes Sordariomycetes and Saccharomycetes accounted for 89.1% of all ascomycete sequences (Figure S7b); both of these have been observed as contaminants in water lines and are unlikely to comprise significant biomass in a brackish pond (Tischner et al. 2021; Mesquita-Rocha et al. 2013). Other common phyla included Holozoa, Ciliophora, and Diatomea, with animal phyla including Cnidaria and Gastrotricha in lower abundance (Figure 3b). Profiling of the 12S rRNA gene suggested the presence of fish associated with brackish water and diadromous migrators (Figure 3c). The most abundant of these was the mummichog *Fundulus heteroclitus*, which had an average abundance of 47.2% across the two samples, followed by sequences classified only to the level Teleostomi (21.4%), a group that includes all bony fish and tetrapods. Other species observed include the Atlantic silverside *Menidia menidia* (19.9%), the American eel *Anguilla rostrata* and unclassified *Anguilla* (9.8% total), and Atlantic herring *Clupea harengus* (1.2%).

**Figure 3.**
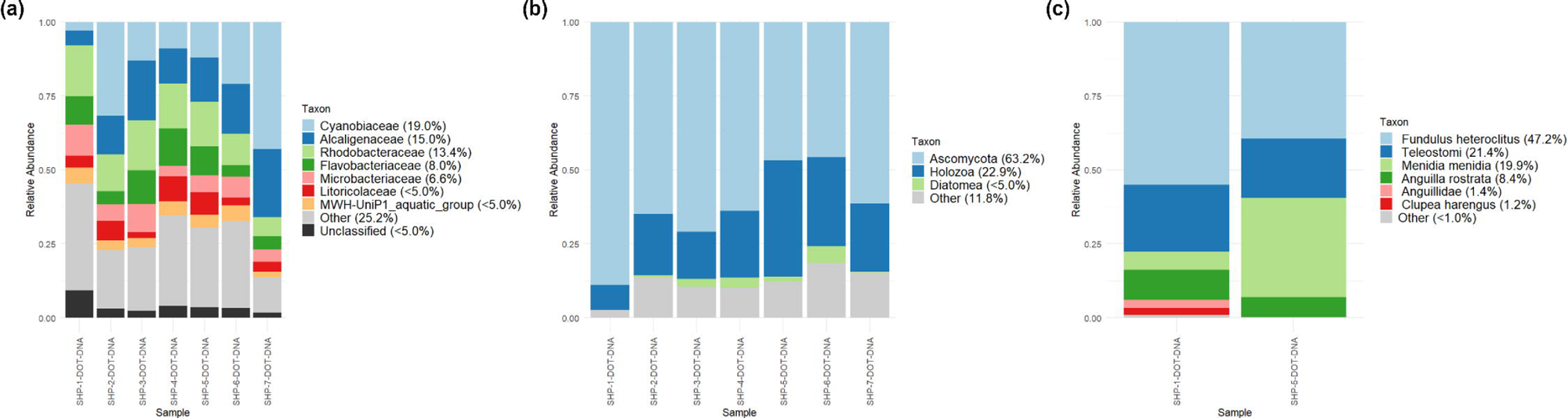
Distributions of taxa based on analysis of marker genes in Sheeps Head Pond. (a) 16S profiles, family level, minimum abundance ≥ 5% in at least one sample. (b) 18S profiles, phylum level, minimum abundance ≥ 5% in at least one sample. (c) 12S profiles, species level, minimum abundance ≥ 1% in at least one sample.

### Grand Lake

Shubenacadie Grand Lake (Mi’kmaw: Kji-qospem, “grand lake”; referred to here simply as Grand Lake) is a freshwater lake in Nova Scotia, approximately 35 km north of Halifax. Grand Lake is a popular fishing location and contains fish including the chain pickerel *Esox niger* and the smallmouth bass *Micropterus dolomieu*. Cyanobacterial blooms are a significant concern; two dogs died in a 2021 incident which was likely due to ingestion of the cyanotoxins produced by the benthic cyanobacterium *Microcoleus*. A subsequent two-year sampling program identified *Microcoleus* at several locations in a stream feeding Grand Lake and proposed that mats may have been transported to the lake shore after a heavy rainfall (Johnston et al. 2024).

Distribution patterns of taxonomic families inferred from the 16S marker showed a striking pattern, with 21 taxa observed in the two RNA samples but missing from all DNA samples including the paired time points 5 and 8 (Figure S8). An additional 16 families were found in one RNA sample and missing from all other DNA and RNA samples, and other common profiles included one or both RNA samples. Sixteen families were present in all DNA and RNA samples, or missing only from sample 6, which had only 188 sequences that passed quality control. Most taxa inferred from the 18S and 12S markers were ubiquitous. Consistent with the high diversity of RNA sequences seen in the UpSet plots, the two DOT and two Niskin samples were clustered in the heatmaps, while DNA samples were interleaved. Many families were ubiquitous or nearly so, including the cyanobacterial family Cyanobiaceae, which are generally found in the water column and not involved with harmful algal blooms (Fukushima et al. 2017; Schallenberg et al. 2021) (Figure S9). 18S profiling split the DOT and Niskin samples completely, with DOT samples containing more Ascomycota and fewer samples with Rotifera, Centrohelida, and Diatomea. Comparing DNA and RNA abundance (Figure S10) showed a negative CLR difference (i.e., RNA CLR values < DNA CLR values) for all taxa considered with an absolute CLR difference of 1.0 or greater. Although this may seem counterintuitive given the increased diversity of the RNA samples, the CLR correction penalizes this diversity when taxon frequencies are divided by the harmonic mean of the sample. The most prominent underrepresentations were Actinobacteriota in the 16S profiles, and Mollusca in the 18S profiles.

Taxonomic profiling showed patterns broadly consistent with a freshwater lake. The 16S profiles were dominated by the families Chitinophagaceae, Comamonadaceae, and Sporicthyaceae, all common constituents of freshwater lakes (Vaughn and Jackson 2024; Newton and McLellan 2015), with many other freshwater bacterial families identified (Figure 4a). Cyanobiaceae, the lone cyanobacterial family, had an average relative abundance of 3.9%. The 18S profiles were dominated by the cryptomonad algae Cryptophyceae (51.9%) and Ciliophora (Figure 4b). Ascomycota were present at much lower levels than were observed in the SHP samples. The underrepresentation of Ascomycota in the RNA samples (Figure S10) may, however, be suggestive of lower levels of contamination. The 12S profiles showed a range of expected freshwater species (Figure 4c), including native species of shad, stickleback, alewife, American eel, and brown bullhead, the invasive smallmouth bass, and the pike family Esocidae, which includes the chain pickerel and other species. Most species were found at both assessed time points except for the alewife and Esocidae.

**Figure 4.**
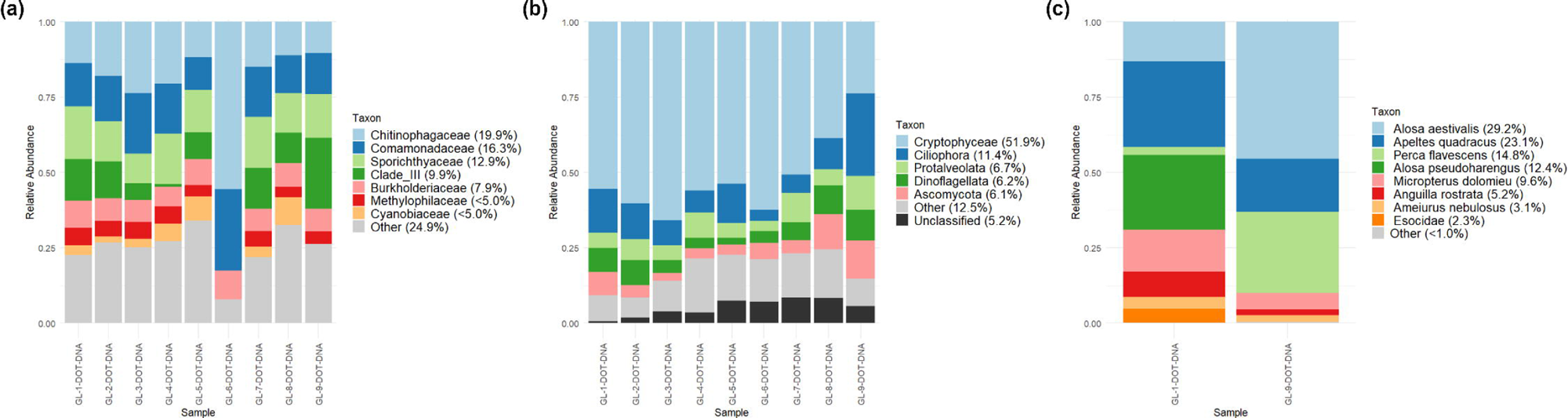
Distributions of taxa based on analysis of marker genes in Grand Lake. (a) 16S profiles, family level, minimum abundance ≥ 5% in at least one sample. (b) 18S profiles, phylum level, minimum abundance ≥ 5% in at least one sample. (c) 12S profiles, species level, minimum abundance ≥ 1% in at least one sample.

### Lunenburg Harbour

Lunenburg, Nova Scotia, (Mi’kmaw: E’se’katik, “place of clams”) is a town with 2400 inhabitants on the South Shore of Nova Scotia. Its harbour has been a focal point of shipbuilding, fish processing, and aquaculture, as well as tourism and recreation. Wastewater contamination is a serious concern in Lunenburg Harbour, and the town has developed an extensive program of upgrades and outfall relocations (Town of Lunenburg 2025). As with many harbours, shipping and other industries have led to significant impacts from invasive species that can replace indigenous species, foul moorings and water intakes, and impact use of the harbour. Such species include the European green crab (*Carcinus maenas*), several species of invasive and established tunicates such as the European sea squirt *Ascidiella aspersa* and compound sea squirt *Diplosoma listerianum* (Moore et al. 2014), and invasive red algae *Dasysiphonia (Heterosiphonia) japonica* (Savoie and Saunders 2013).

Twenty-four prokaryotic families were observed across all DNA and RNA samples, with the remaining common distribution patterns comprising taxa unique to a single sample (Figure S11a). Eleven taxa were found exclusively in one or both RNA profiles. Seventeen phyla inferred from 18S sequences were present in all DNA and RNA samples, with six restricted to either RNA time-point 4, or both RNA profiles (Figure S11b). Seven of eleven species inferred from 12S were present in both profiled samples (Figure S11c). Heatmaps confirmed the presence of most taxa across most or all DOT and Niskin samples at a lower abundance threshold, with all but one bacterial family present in at least 14 of 18 samples (Figure S12a). The 18S phyla followed a similar pattern, with all but five phyla identified in 14/18 samples (Figure S12b). Despite these patterns that were consistent across both DOT and Niskin samples, the two samplers formed separate clusters, likely due to differences in relative abundance. For example, the bacterial families Balneolaceae and SAR116 were overrepresented in DOT samples, as were the eukaryotic phyla Ascomycota, Tunicata, and Picozoa. Several taxa showed an elevated level of RNA abundance relative to DNA: most of these were relatively rare, but several genera such as *Provetella* and *Acinetobacter* had elevated expression levels relative to their relative abundance in the two sequenced samples (Figure S13).

The most-common bacterial families were all marine associated (although not necessarily exclusively) including Flavobacteriaceae, Rhodobacteraceae, Alteromonadaceae, Saprospiraceae, “Clade 1”, and Nitroncolaceae (Figure 5a). We further examined the relative abundance of seven bacterial families often associated with the human gut and other animals: Bacteroidaceae (phylum Bacteroidota); Christensenellaceae, Lachnospiraceae, Enterococcaceae, and Ruminococcaceae (phylum Bacillota); and Enterobacteriaceae (phylum Pseudomonadota) (Figure 6a). The maximum abundance of any of these groups was 0.326% (Prevotellaceae at time-point 7), with no other observation > 0.2%. Enterococcaceae, previously identified as the key fecal contaminant in Lunenburg Harbour, was not detected at any time point. The 18S profiles showed a range of phyla found in both marine and freshwater habitats, including diatoms, cryptophytes, and chlorophyte algae, as well as marine-specific lineages including haptophyte algae and Picozoa (Figure 5b). Tunicates were detected in all samples. 12S profiling showed a range of known marine and brackish taxa, including sculpins (Cottidae), which were classified only to the family level; the winter flounder *Pseudopleuronectes americanus*, and American eel *Anguilla rostrata*, which were all found in both samples; and other species including the alewife *Alosa pseudoharengus*, rock gunnel *Pholis gunnellus*, and Atlantic menhaden *Brevoortia tyrannus*, which were found at lower abundance (Figure 5c).

**Figure 5.**
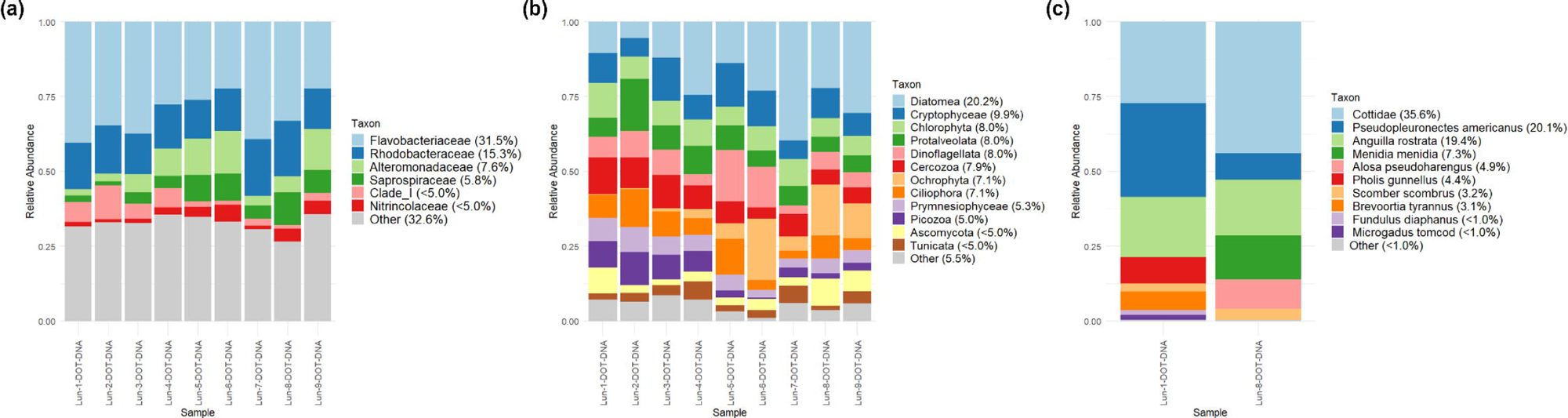
Distributions of taxa based on analysis of marker genes in Lunenburg Harbour. (a) 16S profiles, family level, minimum abundance ≥ 5% in at least one sample. (b) 18S profiles, phylum level, minimum abundance ≥ 5% in at least one sample. (c) 12S profiles, species level, minimum abundance ≥ 1% in at least one sample.

**Figure 6.**
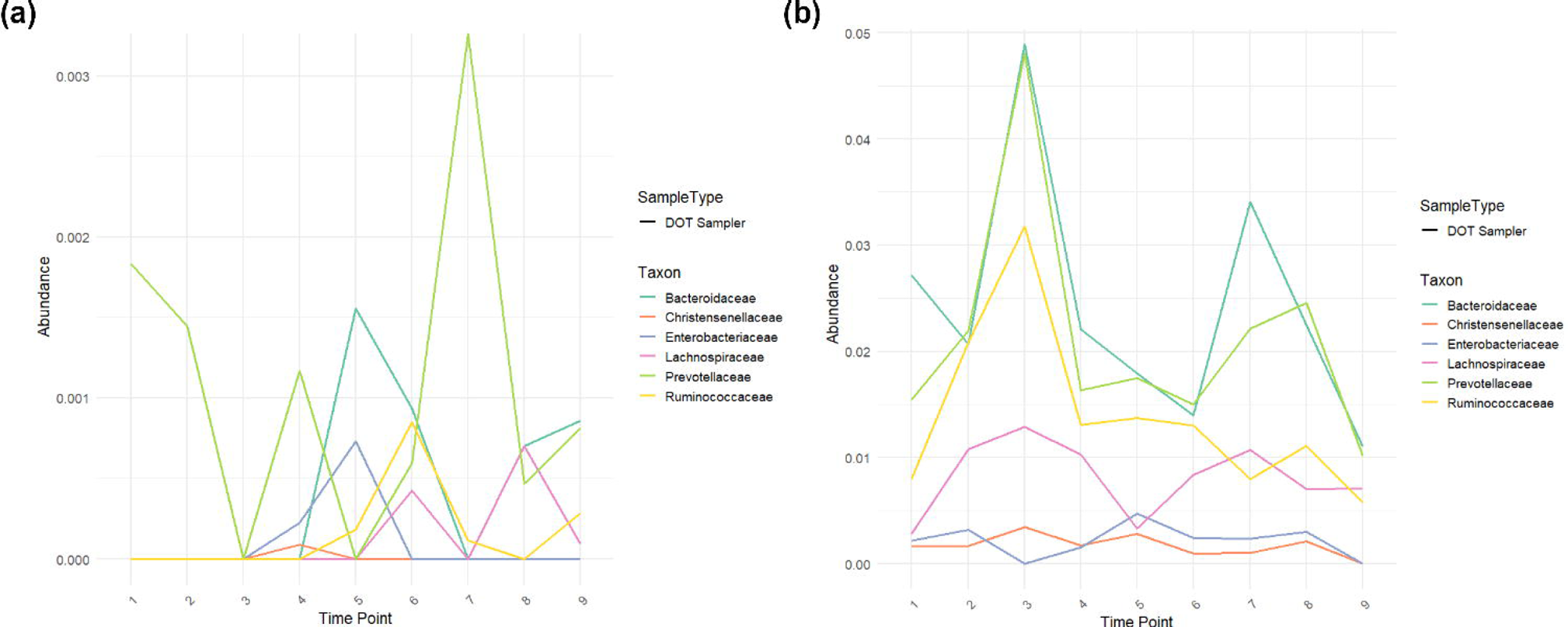
Relative abundance of six bacterial families across nine time points in samples collected from Lunenburg Harbour. (a) and COVE Jetty (b). A seventh family, Enterococcaceae, was absent from all collected samples.

### COVE Jetty, Halifax Harbour

Halifax, Nova Scotia (Mi’kmaw: Kjipuktuk, “great harbour”), is the largest city on the east coast of Canada, with approximately 500,000 inhabitants. Halifax Harbour is a working harbour that includes shipyards, power stations, naval yards, multiple container ports, and shipbuilding, as well as recreation and tourism. For centuries, untreated wastewater was dumped into the harbour, leading to poor water quality and accumulation of toxins in sediment (Buckley and Winters 1992). Wastewater treatment upgrades in the late 2000s resulted in improved water quality and lower microbial loads, but in times of high rainfall, sewage is still pumped directly into the harbour. On July 21-22, 2023, Nova Scotia experienced a massive rainfall event with up to 250 mm falling in parts of the Halifax Regional Municipality in a 24-hour period, which led to catastrophic flooding (CTV News Atlantic 2023). On July 22, we deployed a DOT sampler near a sewage outflow to assess the potential increases in human-associated bacteria.

Similar to Lun, taxonomic groups inferred from CJ samples tended to be either ubiquitous across all DNA and RNA profiles, or restricted to a single sample (Figure S14). Thirty-one prokaryotic families were found in all DNA and RNA samples, with the next four profiles by abundance corresponding to taxa present only in a single profile (32 taxa total). Fifteen 18S phyla were ubiquitous, with no other presence/absence profile containing more than two taxa. Similarly, nine species identified using 12S were found in both samples, with the remaining two found only in the time-point 9 sample. Heatmaps of taxon presence/absence showed even more consistency than they did for Lun, likely due in part to the absence of Niskin samples (Figure S15). Two families typically associated with the human microbiome, Ruminococcaceae and Lachnospiraceae, were present with abundance ≥ 1% in six and four profiles, respectively, although both were absent from the RNA profiles. The animal phyla Tunicata, Mollusca, and Holozoa were seen consistently across all samples, apart from one RNA profile in which Mollusca were not observed (Figure S15b). No prokaryotic families showed a CLR difference ≥ 1% between paired DNA and RNA samples. In contrast, many eukaryotic phyla showed overrepresentation above this threshold, while only Tunicata were underrepresented (Figure S16).

Consistent with the patterns shown in the PCoA plots (Figure 2), the dominant bacterial families in CJ were similar to those found in Lunenburg Harbour, including Flavobacteriaceae, Rhodobacteraceae, Alteromonadaceae, Saprospiraceae, and “Clade 1” present in the same pattern of decreasing order (Figure 7a). However, three other families were found in at least one CJ sample with a relative abundance ≥ 5%: the NS9 marine group, Comamonadaceae, and Balneolaceae. We further examined the relative abundance of seven bacterial families often associated with the human gut and other animals: Bacteroidaceae (phylum Bacteroidota); Christensenellaceae, Lachnospiraceae, Enterococcaceae, and Ruminococcaceae (phylum Bacillota); and Enterobacteriaceae (phylum Pseudomonadota) (Figure 6b). None of these groups was observed with relative abundance > 0.326% in Lun, so a higher abundance in Halifax Harbour could be consistent with the release of a greater volume of untreated sewage. Four of these six groups were present at a relative abundance > 1% in at least one CJ sample; Bacteroidaceae was the eighth most-abundant bacterial family overall with an average abundance of 2.4% (Figure 6b), and the only family not among the top ten most-abundant families in Lunenburg Harbour. Five of the six families peaked in relative abundance at time-point 3 (12:02 AM, July 23), with Bacteroidaceae and Prevotellaceae at an abundance of nearly 5%, Ruminococcacae above 3%, and Lachnospiraceae above 1%. 18S phylum-level profiles were similar to those found in Lun, albeit with different average relative abundance values: for example, Cryptophyceae (CJ = 26.8%, Lun = 9.9%), Prymnesiophyceae (CJ = 14.6%, Lun = 5.3%), and Chlorophyta (CJ = 14.3%, Lun = 8.0%) (Figure 7b). Fewer species were present with relative abundance ≥ 1% in the 12S profiles: the Atlantic mackerel *Scomber scombrus* (CJ: 38.2%, Lun: 3.2%), the alewife *Alosa pseudoharengus* (CJ: 24.3%, Lun: 4.9%), the winter flounder *Pseudopleuronectes americanus* (CJ: 18.6%, Lun: 20.1%), and the rock gunnel *Pholis gunnellus* (CJ: 8.1%, Lun: 4.4%). Additionally, 10.1% of CJ 12S sequences were assigned to the order Zoarcales, which may also correspond to gunnels.

**Figure 7.**
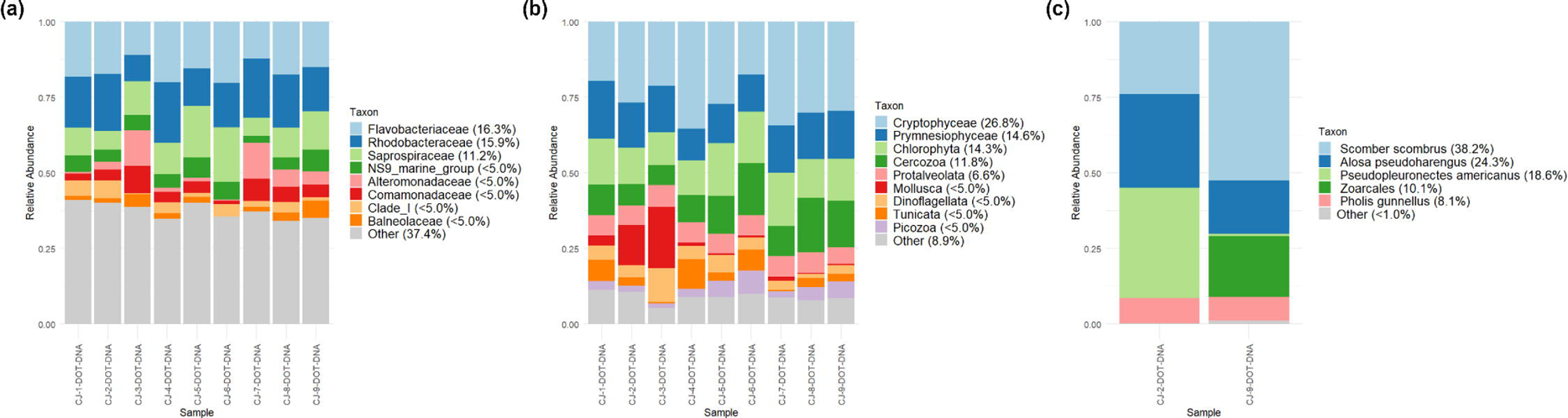
Distributions of taxa based on analysis of marker genes in COVE Jetty. (a) 16S profiles, family level, minimum abundance ≥ 5% in at least one sample. (b) 18S profiles, phylum level, minimum abundance ≥ 5% in at least one sample. (c) 12S profiles, species level, minimum abundance ≥ 1% in at least one sample.

Metagenomic sequencing was successful for all samples, with an average of 16,680,906 reads per sample. The generated assemblies produced N50s between 4749 and 8350, with the longest contigs ranging in size between 129,130 and 415,927 nt (Table S2). The longest contigs mapped to several different genera, including *Planktomarina* (family Rhodobacteraceae, samples 1, 2, 3, and 9); *Pseudomonas* (family Pseudomonadaceae, samples 4 and 8); *Flavobacterium* (family Flavobacteriaceae, samples 5 and 6); and *Alteromonas* (family Alteramonadaceae, sample 7). All but *Pseudomonas* were among the five most-abundant families from the 16S profiles; by contrast, *Pseudomonas* was completely absent from the CJ 16S profiles, suggesting the Kraken assignments may be a misclassification of a contig of heterogeneous origin. Other contigs of length > 100,000 in samples 4 and 8 mapped to families Rhodobacteraceae, Flavobacteriaceae, and Alteromonadaceae. There was considerable disagreement in the relative abundance of bacterial families between the 16S samples and Kraken predictions (Figure S16). For example, Pelagibacteraceae were absent from the 16S profiles but were the most abundant in the metagenomic sequence read and contig classifications among the 20 families shown in Figure S16. Conversely, several families from Proteobacteria and Bacteriodota and OM190, the lone family from Planctomycetota, were found in the 16S but not the metagenomic samples.

Metagenomic functional profiling of key GO pathways yielded normalized estimates of relative pathway abundance. Key nutrient-cycling pathways including photosynthesis, sulfur, phosphate, nitrogen, and carbon metabolism (Figure 8), none of which showed an increasing, decreasing, or cyclic trend. We also used RGI with the CARD database to assess the presence of putative antimicrobial-resistance genes (ARGs) from the assembled contigs. A maximum of 15 Perfect and Strict ARGs were predicted in any sample, with glycopeptide and disinfecting agents comprising 78.8% of all predicted genes (Table S3-S6). However, percent identities for most resistance classes were generally low (<50%), apart from five genes predicted to confer resistance to macrolides and streptogramins (100% identity to reference sequence), which mapped to the family Enterobacteriaceae, and four genes labeled “fluoroquinolone; diaminopyrimidine; phenicol” (86.7% identity) predicted to be present in Vibrionaceae. The apparent rarity and poor conservation of ARGs is consistent with the taxonomic profile of the samples, where most reference CARD genes are from pathogenic organisms (e.g., Enterobacteriaceae), while the CJ samples are dominated by classes such as Alphaproteobacteria that are poorly represented or absent from validated reference databases.

**Figure 8.**
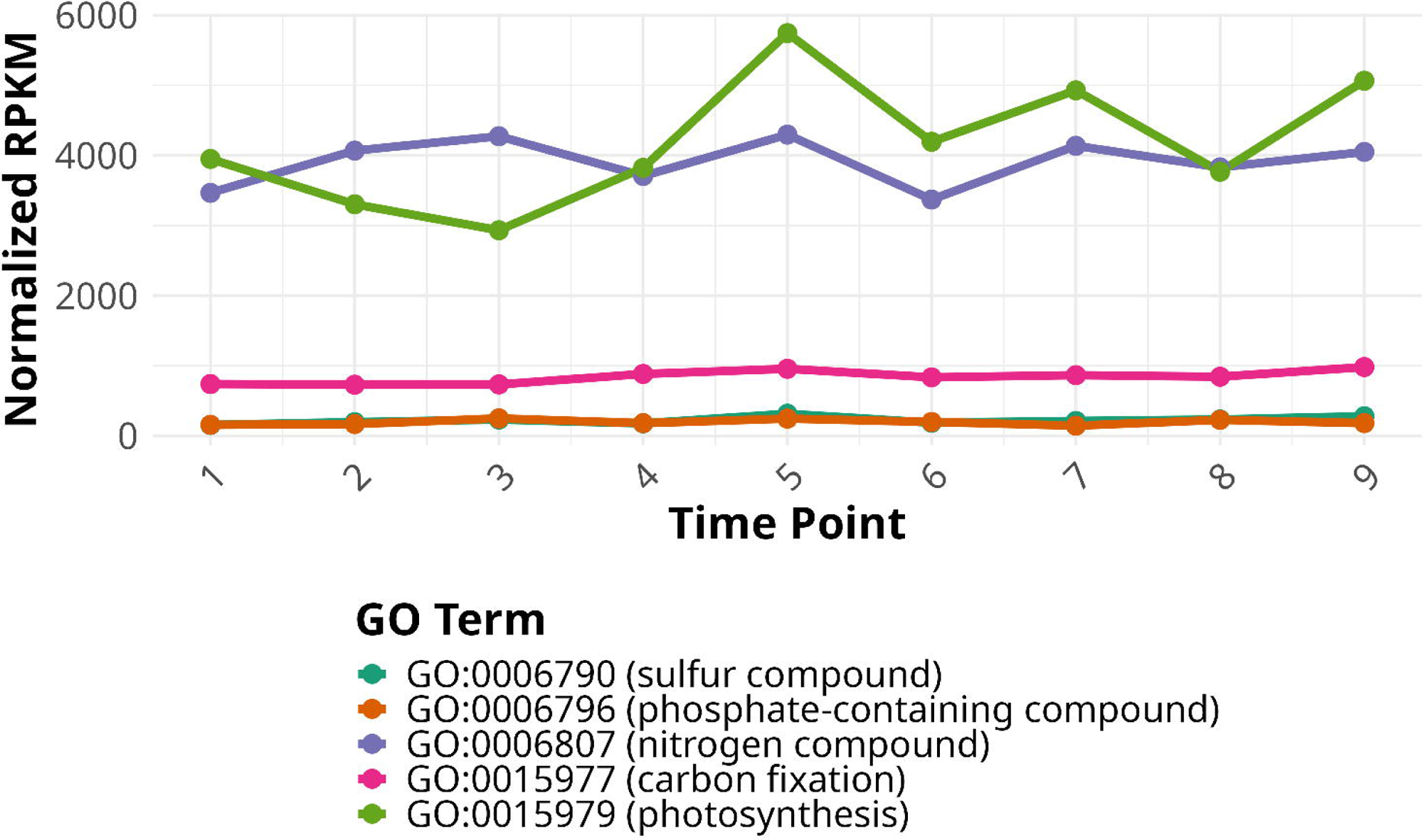
Normalized read counts of genes associated with key biogeochemical cycling pathways and photosynthesis, inferred using HUMAnN3.

## Discussion

Our study provides a novel demonstration of the utility and versatility of a robust autonomous sampler across a range of aquatic environments. We assessed the DOT sampler’s capacity to preserve and recover high-quality DNA, and compared its performance to conventional sampling methods using multiple genetic markers. The successful deployment at the COVE Jetty, along with previous longer-term deployments (Van Wyngaarden et al. 2024), demonstrates its utility in collecting samples in challenging conditions, remote locations, and at otherwise inconvenient times (e.g., overnight). By integrating eDNA and eRNA metabarcoding together with metagenomic profiling, we also highlight the unique temporal and biological insights afforded by the combination of eDNA and eRNA analyses. Our findings demonstrate both the diversity and temporal dynamics of these systems, and underscore the potential for autonomous molecular monitoring tools to enhance long-term monitoring of aquatic ecosystems. These advances in autonomous molecular monitoring directly respond to the growing need for scalable biodiversity assessment tools that can support conservation initiatives and effectively measure ecosystem change. As global targets drive the expansion of protected areas and new conservation measures, innovative tools for biodiversity monitoring, such as autonomous samplers, will be essential for evaluating effectiveness and guiding management across diverse and dynamic aquatic environments.

### Sampling

We successfully used the DOT eDNA sampler to profile the biodiversity of different groups of organisms with a range of molecular and bioinformatic techniques. Despite the small volumes obtained from both DOT and Niskin samplers in this study, sequence recovery in nearly all samples was >4000 reads, with the AllPrep® extractions yielding especially high read counts. Crucially, most taxa found in each habitat were found in all samples, or a single sample only (Figure S4, S8, S11, S14). Our results are consistent with Van Wyngaarden et al. (2024) who detected the most common marine fishes and invertebrates in Halifax Harbour from only 125mL of seawater. Together with our current study, these results highlight how even small volumes of water can yield significant DNA and RNA concentrations for metabarcoding and metagenomics, at least in shallow waters rich in biomass and species diversity. Future tests will encompass the deep sea, where eDNA concentrations are generally lower than shallow or surface waters, and greater volumes of water may be necessary to characterize fish and invertebrate communities (e.g., (McClenaghan 2020)). Quantitative PCR or digital droplet PCR from preserved samples or custom metabarcoding primers will be necessary if expected DNA abundance is extremely low or if closely related species must be reliably differentiated (Wilcox et al. 2013; Nathan et al. 2014). Ordination plots (Figure 2) demonstrated a high degree of concordance between the DOT and Niskin samples, even for the SHP samples where the DOT sampler showed evidence of contamination by ascomycete fungi in the 18S profiles. There were differences in the diversity estimates and taxonomic profiles generated by the two samplers, but these trends were not systematic and both samplers identified taxa that were not predicted by samples from the other. This finding is consistent with patterns reported previously in a more-limited trial (Hendricks et al. 2023).

### Contrasting eDNA and eRNA

We obtained high eDNA and cDNA concentrations from the AllPrep® kits (average sequencing depth = 83,541 reads / sample). Although sample sizes were too small for robust statistical comparisons, we were able to demonstrate substantial log-ratio differences for taxa with varying degrees of abundance. Further, taxonomic profiles inferred from eRNA were often more similar to each other than they were to the matched eDNA samples in ordination plots and clustered heatmaps. Many taxa were observed only in eRNA profiles, most notably in Grand Lake, where 21 bacterial families were inferred only from the two eRNA samples (Figure S8). eRNA in our study thus yielded higher species richness, particularly at the microbial level, suggesting it better reflects species composition of active bacteria at the time of sampling. eRNA allows the comparison of taxa based on metabolic activity and can better differentiate between active, less-active, and dead organisms due to different decay rates (Kagzi et al. 2022; Jo et al. 2022); eRNA also has the capability to identify variation on different time scales than eDNA. In lakes and other aquatic environments, eRNA has demonstrated significantly greater true positive rates of fish detections relative to eDNA (Littlefair et al. 2022). While this result might seem counterintuitive given the lesser stability of RNA, several factors may increase the persistence of RNA, including differences in particle association and higher copy numbers of ribosomal RNA versus DNA in metabolically active cells (Torti et al., 2015). Successful preservation and amplification of standard eRNA marker genes suggests the possibility of performing targeted amplification of specific functional marker genes, and metatranscriptomic analysis to assess the expressed functional profile of the microbiome (Shakya et al. 2019; Yates et al. 2021).

### Location-specific insights

We used the DOT and Niskin samples to address general questions about inferred biodiversity (e.g., marine vs. freshwater taxa, invasive species) and targeted questions for each site. Bacterial taxa observed in Grand Lake such as Chitinophagaceae, Comamonadaceae, and Sporichtyaceae were primarily freshwater associated. In contrast, the majority of bacterial taxa in Lunenburg Harbour and at the COVE Jetty were marine associated, including the families Flavobacteriaceae, Rhodobacteraceae, and Alteromonadaceae. Additionally, the COVE Jetty had lower-abundance signatures of potential runoff and wastewater-associated bacteria such as Comamonadaceae and the probable human-associated families shown in Figure 6. The seven human microbiome-associated families examined were approximately ten times higher in abundance in the COVE Jetty samples than in Lunenburg Harbour, which may reflect improved wastewater treatment in Lunenburg and the sewage discharge that followed the major rainfall event in Halifax, but thorough tests with baselines will be necessary to demonstrate this more conclusively. The brackish Sheeps Head Pond contained a mix of bacteria associated with marine, fresh, and brackish waters, including Cyanobiaceae, Rhodobactereaceae, and Litoricolaceae, with small but detectable levels of Desulfobacterota.

Eukaryotic profiles from 18S, summarized at the phylum level, were also consistent with aquatic habitat types: Cryptophyceae, Ciliophora, and Protalveolata in the freshwater lake; Diatomea, Cryptophyceae, Chlorophyta and other marine taxa in Lunenburg Harbour; and the COVE Jetty showed a mix of the Lunenburg signature with a higher proportion of Cryptophyceae, Cercozoa, and Protalveolata. Animal phyla were present but less abundant in the 18S data, including Holozoa, Arthropoda, Rotifera, and other taxa in Sheeps Head Pond (Figure S5b), Holozoa, Porifera, Mollusca, and Rotifera in Grand Lake (Figure S9b), and Mollusca and Tunicata in both harbours, with Arthropoda and Annelida additionally present in Lunenburg Harbour (Figure S12b, Figure S15b). 12S profiling detected fish signatures that were too uncommon to be reflected in the 18S data, including the coastal-associated mummichog and estuarine Atlantic silverside in Sheeps Head Pond; freshwater species including yellow perch, smallmouth bass, and pike in Grand Lake; and mackerel, alewife, and winter flounder in both harbours, along with several other species identified in Lunenburg Harbour. These fish species are consistent with those detected by Van Wyngaarden et al. (2024) in Halifax Harbour in fall 2023.

Metagenomic analysis of COVE Jetty samples identified key nutrient-cycling pathways at similar relative abundances across the nine time points sampled, despite changes in the relative abundance of probable human-associated taxa. Taxonomic profiling of metagenomic versus 16S sequence data was highly inconsistent (Figure S17), with many abundant taxa inferred from 16S showing lower abundance or complete absence from the metagenomic predictions, while the family Pelagibacteraceae showed the opposite pattern. This inconsistency is well documented (Jovel et al. 2016; Rausch et al. 2019; Cordier et al. 2021; Zhao et al. 2023; Kleikamp et al. 2023) and arises due to reference-database incompatibilities (for example, Pelagibacteraceae is absent from the SILVA taxonomy), PCR primer biases, and inconsistencies between the nucleotide patterns in 16S versus whole-genome sequences that are used for classification. These differences highlight the need for careful interpretation of taxonomic profiles. The AMR analysis (Table S3-S6) identified few plausible genes but illustrates the ability to profile specific genes of interest from preserved samples.

### Further development

The filter-cassette design of the DOT sampler allows for rapid swapping out of filters in the field, with the additional benefit of minimizing re-engineering to accommodate a variety of different filter types. The end cap and filter-cassette holder have recently been modified to accommodate larger filters, such as sealed single-use capsule filters, which are 50 mm in outer diameter and internally have a filter stack comprised of a 45 mm diameter 5 μm glass fibre prefilter and a 45 mm diameter 0.8 μm PES membrane (Luy et al. 2024). While the samples collected in this study were generally 100 mL or less, the 45 mm diameter filters will allow the capture of 5 L+ per sampling event, depending on the filter poresize and application. Automated cleaning, flushing of the inlet, and sterilization of the fluid lines ensures that samples captured on the filter membrane/material are representative of the water sample during the time window of capture. The data collected in this work revealed that the cleaning protocol used required refinement as some dead volumes were not flushed with 5% HCl. An updated cleaning protocol has been implemented on the samplers and validated by arms-length third parties, eliminating cross-contamination (Yamahara et al. 2025). The operational range of the sampler has also been extended in new units, with configurations that have depth ratings of 20 m, 200 m, and 2000 m, and a lower-bound operating temperature of –1 to – 2°C, as demonstrated by successful deployments at freezing temperatures in the Southern Ocean. The wetted materials of the eDNA sampler (FKM, PEEK, Teflon, etc.) are compatible with numerous cleaning fluids (e.g., bleach) and preservatives (e.g., DNAshield). However, studies detailing performance using alternative cleaning/preservative combinations would be required before use in long-term deployments. Successful sample preservation also allows for other profiling techniques such as transcriptomics, targeted profiling using qPCR, or sequencing while aboard research vessels or in remote locations.

## Abbreviations

ASV: amplicon sequence variant
eDNA/eRNA: Environmental DNA/RNA
16S/18S/12S: 16S/18S/12S ribosomal RNA gene
SHP: Sheeps Head Pond, Nova Scotia
GL: Shubenacadie Grand Lake, Nova Scotia
Lun: Lunenburg Harbour, Nova Scotia
CJ: COVE Jetty, Halifax Harbour, Nova Scotia
CLR: centered log-ratio transformation
DOT: Dartmouth Ocean Technologies, Inc.

## Author Contributions

RGB, TK, VS, and JL conceived and designed the study. TK, CMM, and IG collected the samples. RGB, JT, SSB, MF, SSNM, NWJ, RRES, VS, and JL analyzed and interpreted the data. All authors contributed to writing of the manuscript.

## Acknowledgments

Funding for this project was provided by Genome Atlantic, the Canada First Research Excellence Fund “Transforming Climate Action” project, and the Natural Sciences and Engineering Research Council of Canada.

## Data Availability

Data will be submitted to a public repository (NCBI or EMBL) prior to resubmission.

