## Supplemental Figures for "Automated eDNA and eRNA Profiling for Biodiversity Monitoring in Marine and Freshwater Ecosystems"

Received: Added at production | Revised: Added at production | Accepted: Added at production
DOI: xxx/xxxx

ARTICLE TYPE

### Supplementary Information for “Automated eDNA and eRNA Profiling for Biodiversity Monitoring in Marine and Freshwater Ecosystems”

 © 2023 Copyright Holder Name 

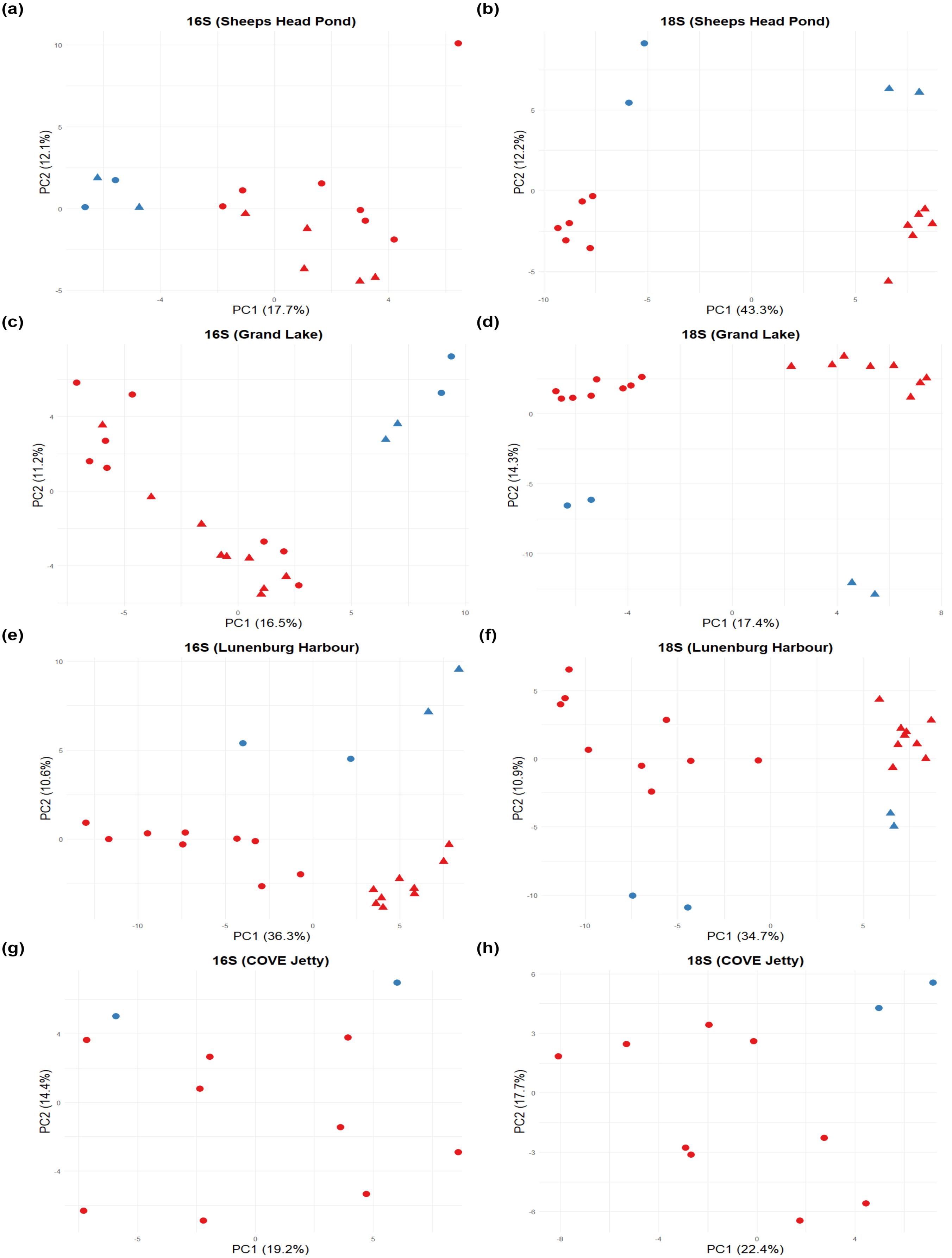


**FIGURE S1** Principal coordinate plots based on the Aitchison distance for 16S (a-d) and 18S (e-h) markers for all locations. Circles indicate DOT samples and triangles Niskin samples. Red = DNA, blue = RNA.

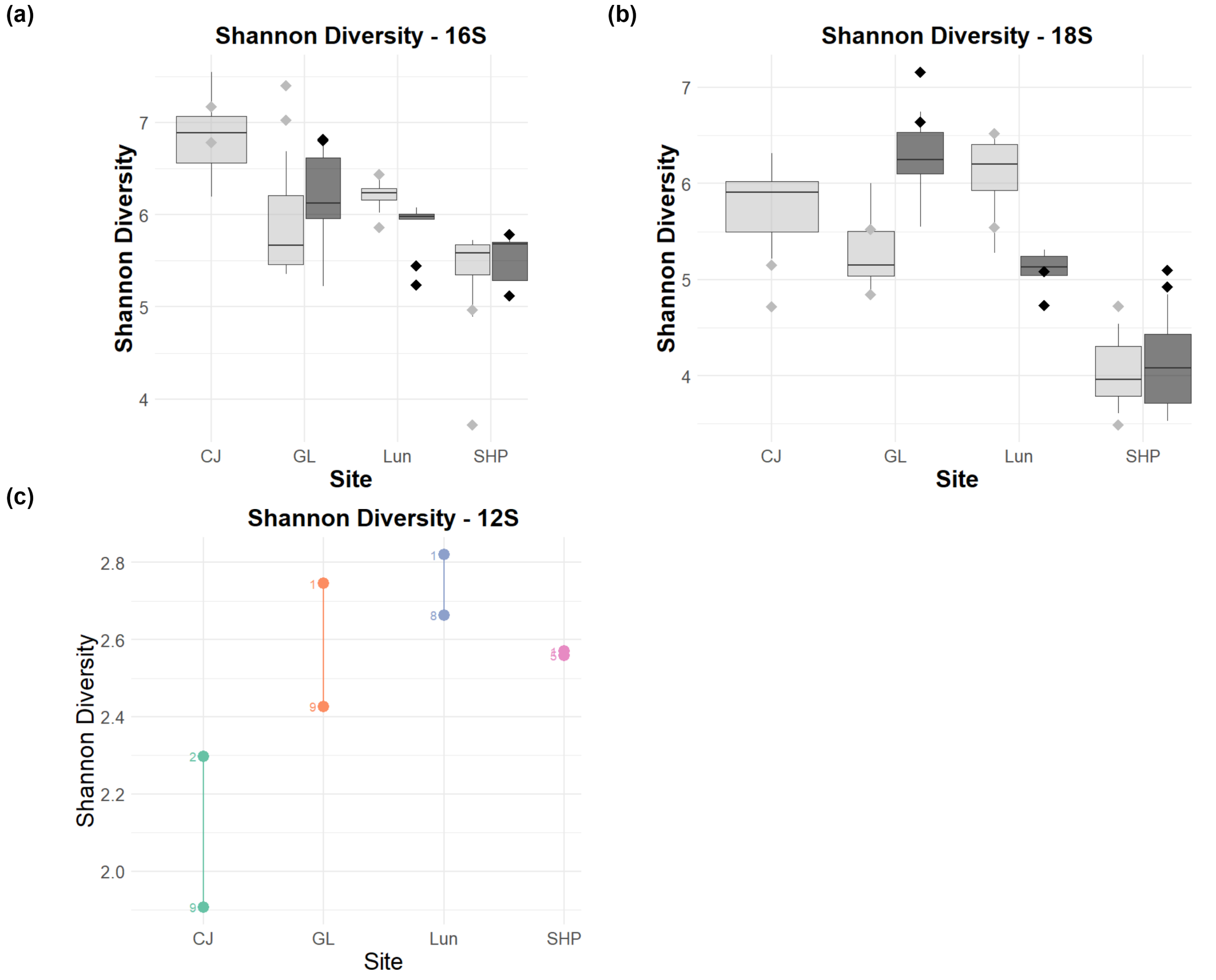


**FIGURE S2** Shannon diversity of samples by marker and location. Diversity scores of DNA samples for 16S (a) and 18S (b) are summarized using boxplots, with the two RNA samples shown with diamonds. Light gray boxes = DOT sampler, dark gray = Niskin sampler. No Niskin samples were collected for CJ. (c) Shannon diversity scores of DOT DNA samples for the 12S marker (two per location, DOT and DNA only). Time points are shown and points are coloured by location.

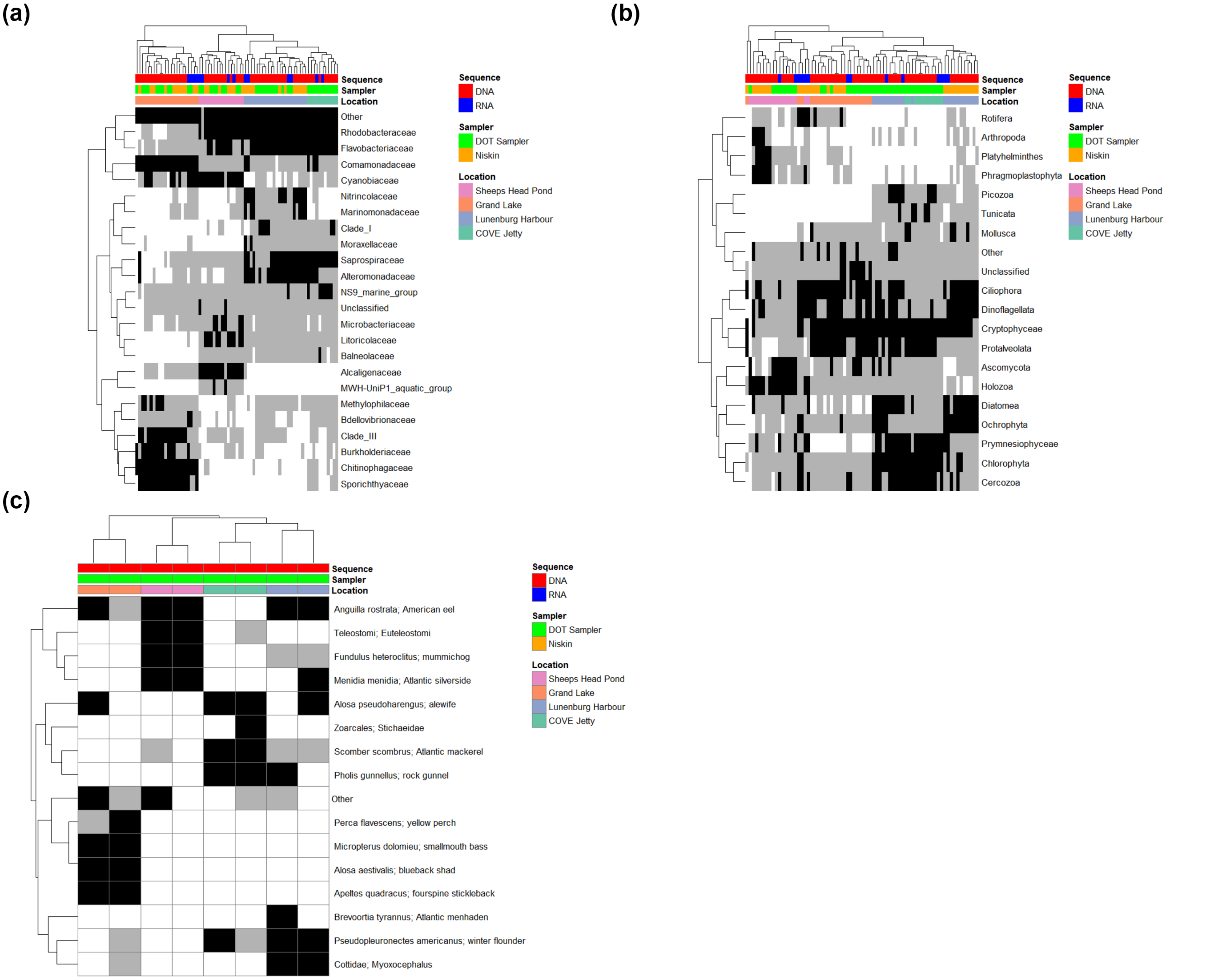


**FIGURE S3** Presence / absence heatmaps for all samples by location, sequence type (DNA vs RNA) and sampler (DOT vs Niskin). All taxa shown are present in at least one sample at a minimum relative abundance of 5%. Samples with taxon abundance above this threshold are shown are indicated with black rectangles, while taxon abundance $\geq$ 0.1% and < 5% are shown in gray.

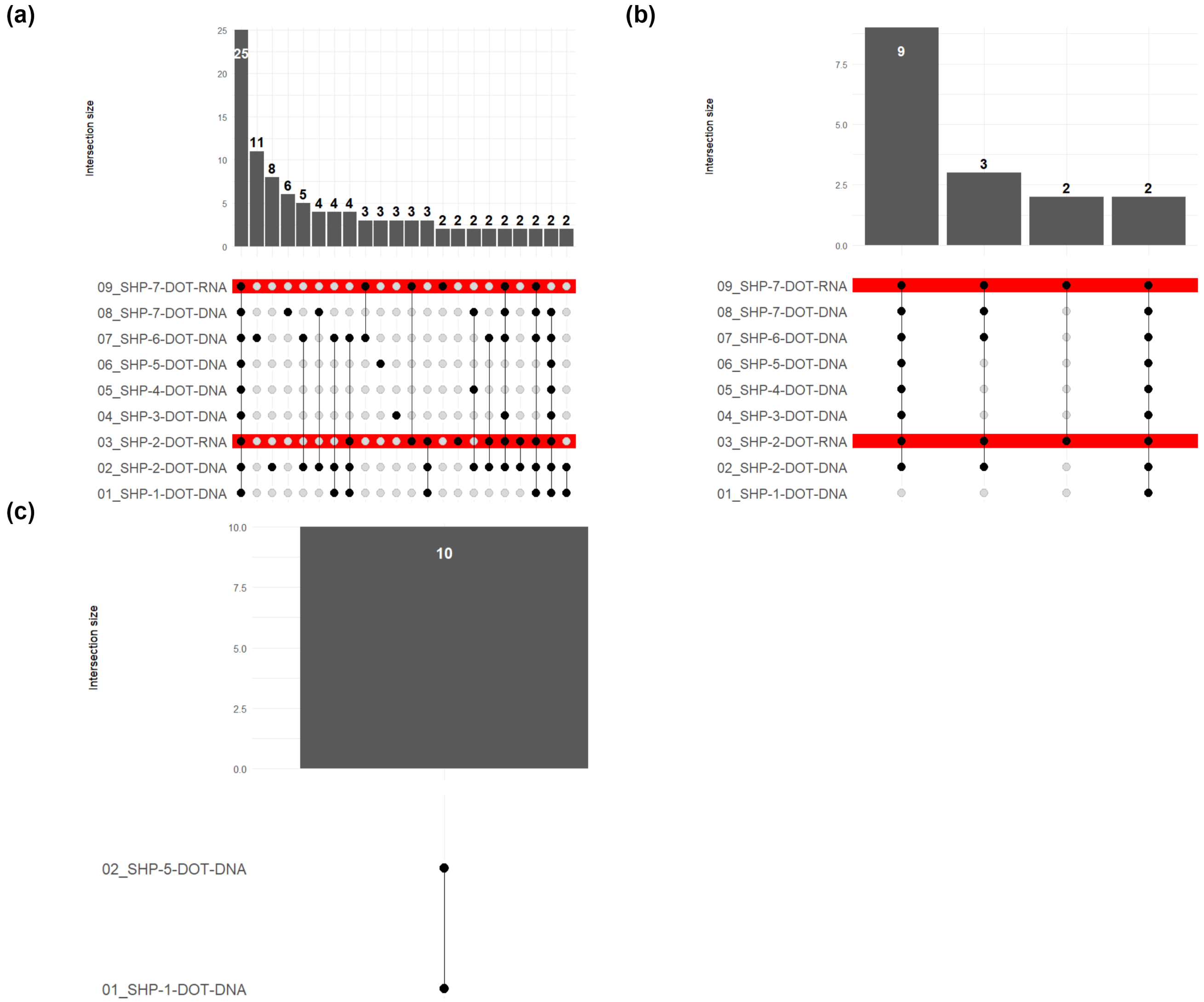


**FIGURE S4** UpSet plots showing distribution of taxa across all DOT DNA and RNA samples for 16S families (a), 18S phyla (b), and 12S species (c) profiles in Sheeps Head Pond. Each column in the plot indicates a unique presence / absence pattern across all samples, with black dots indicating presence and white dots absence. Bars at the top with numbers indicate the number of taxa that exhibit that presence/absence pattern. RNA profiles for 16S and 18S are highlighted in red. Numbers preceding each sample name are for ordering purposes only.

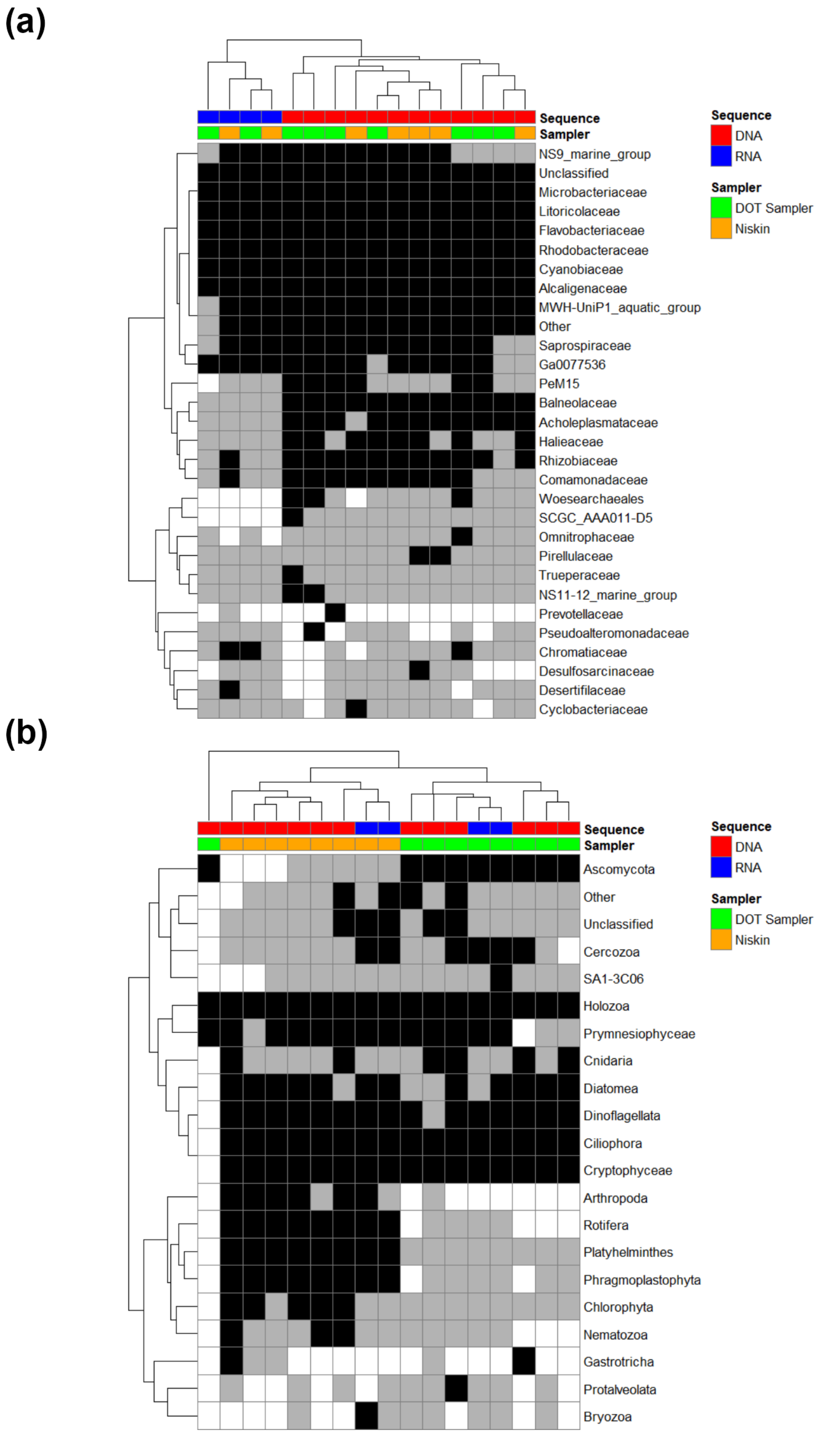


**FIGURE S5** Presence / absence heatmaps for all samples by sequence type and sampler for 16S families (a) and 18S phyla (b) from Sheeps Head Pond. All taxa shown are present in at least one sample at a minimum relative abundance of 1%. Samples with taxon abundance above this threshold are shown are indicated with black rectangles, while samples with taxon abundance less than 1% are shown in gray.

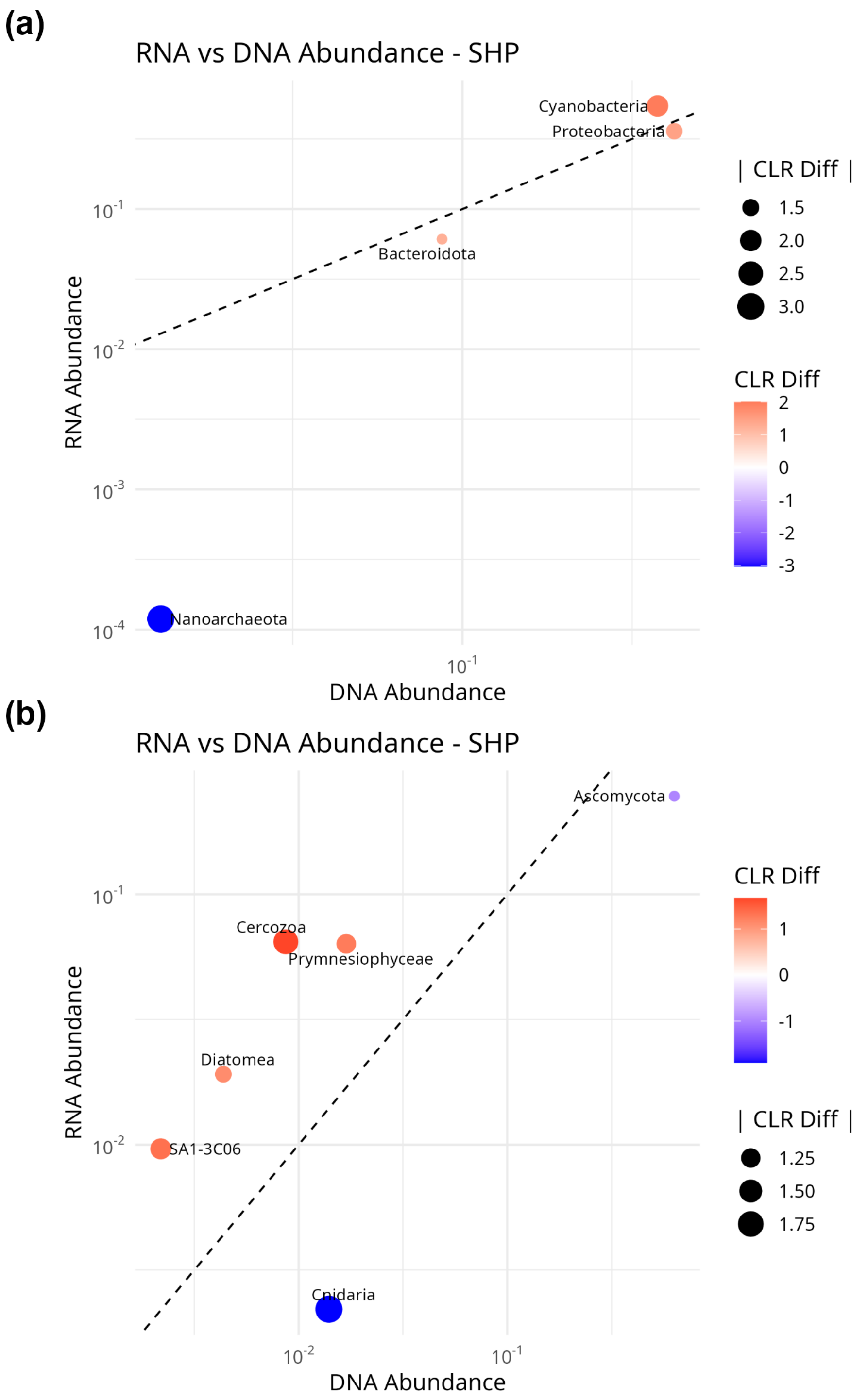


**FIGURE S6** Abundance and centered log-ratio comparisons for paired DNA and RNA samples for 16S families (a) and 18S phyla (b) from Sheeps Head Pond. x- and y-axes show the relative abundance of each taxon in the DNA and RNA samples respectively on a logarithmic scale. The size of each coloured dot indicates the absolute average difference between the CLR scores for paired RNA and DNA samples (see Methods), while the colour scale indicates the magnitude and direction of the difference. Red dots indicate CLR(RNA) > CLR(DNA), while blue dots indicate the opposite pattern. Only paired samples with absolute CLR difference greater than 1.0 are shown.

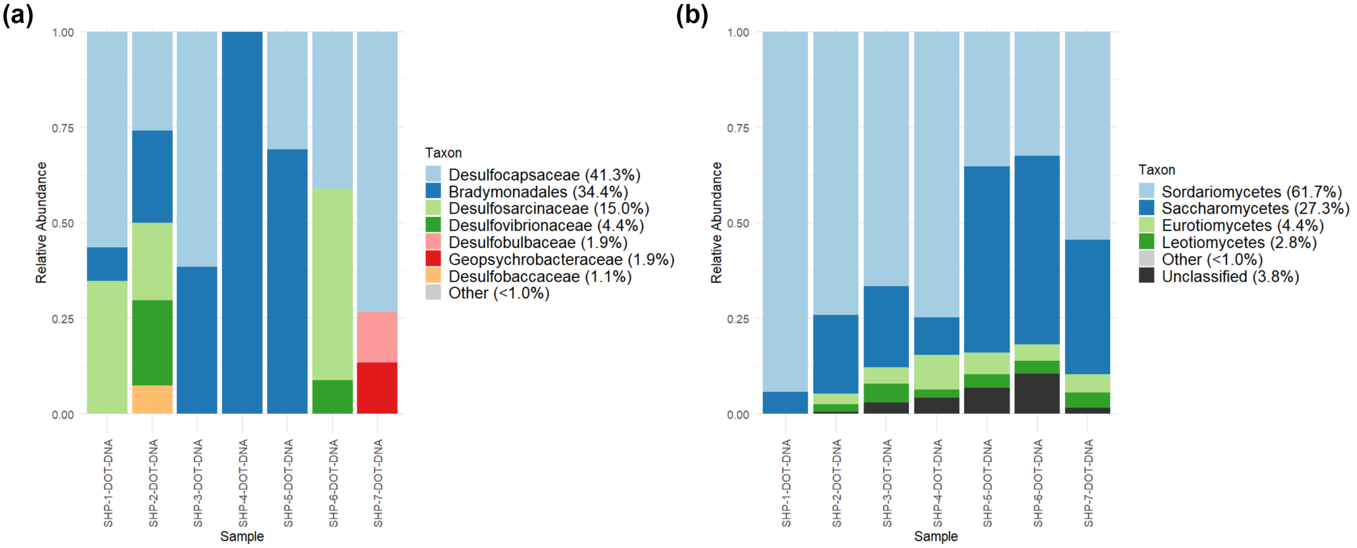


**FIGURE S7** Taxonomic distributions of taxonomic classes within the bacterial phylum Desulfobacterota (a) and the eukaryotic phylum Ascomycota (b). Taxa with abundance $\geq$ 1% in at least one sample are shown.

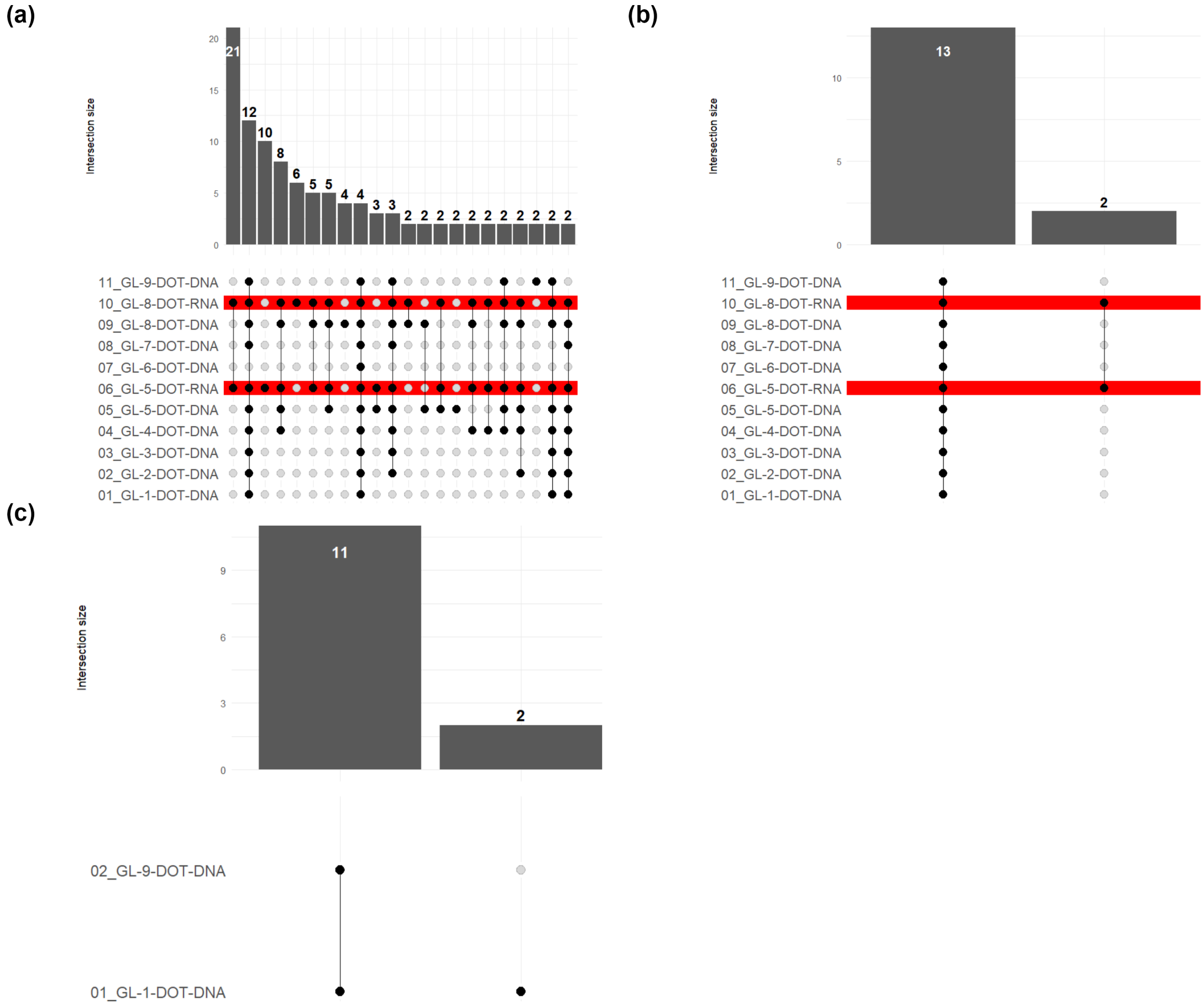


**FIGURE S8** UpSet plots showing distribution of taxa across all DOT DNA and RNA samples for 16S families (a), 18S phyla (b), and 12S species (c) profiles in Grand Lake. Each column in the plot indicates a unique presence / absence pattern across all samples, with black dots indicating presence and white dots absence. Bars at the top with numbers indicate the number of taxa that exhibit that presence/absence pattern. RNA profiles for 16S and 18S are highlighted in red. Numbers preceding each sample name are for ordering purposes only.

(a)

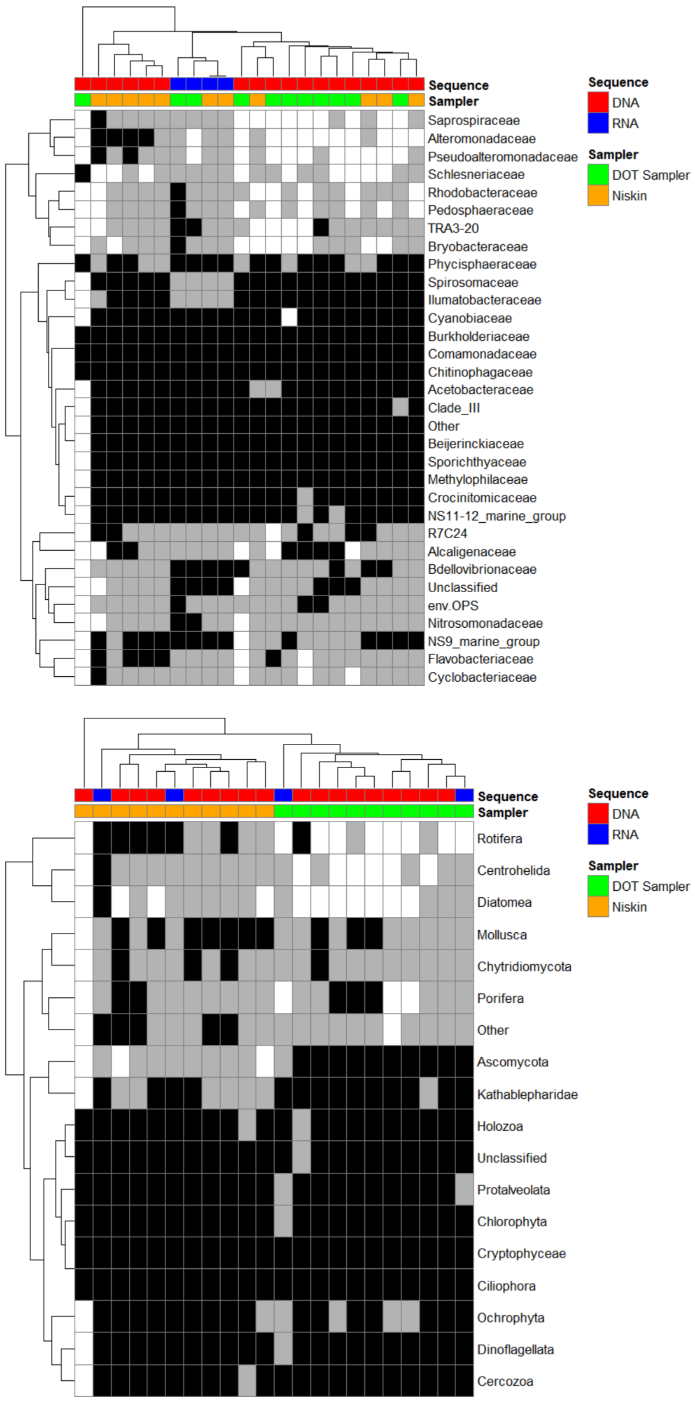


(b)



**FIGURE S9** Presence / absence heatmaps for all samples by sequence type and sampler for 16S families (a) and 18S phyla (b) from Grand Lake. All taxa shown are present in at least one sample at a minimum relative abundance of 1%. Samples with taxon abundance above this threshold are shown are indicated with black rectangles, while samples with taxon abundance less than 1% are shown in gray.

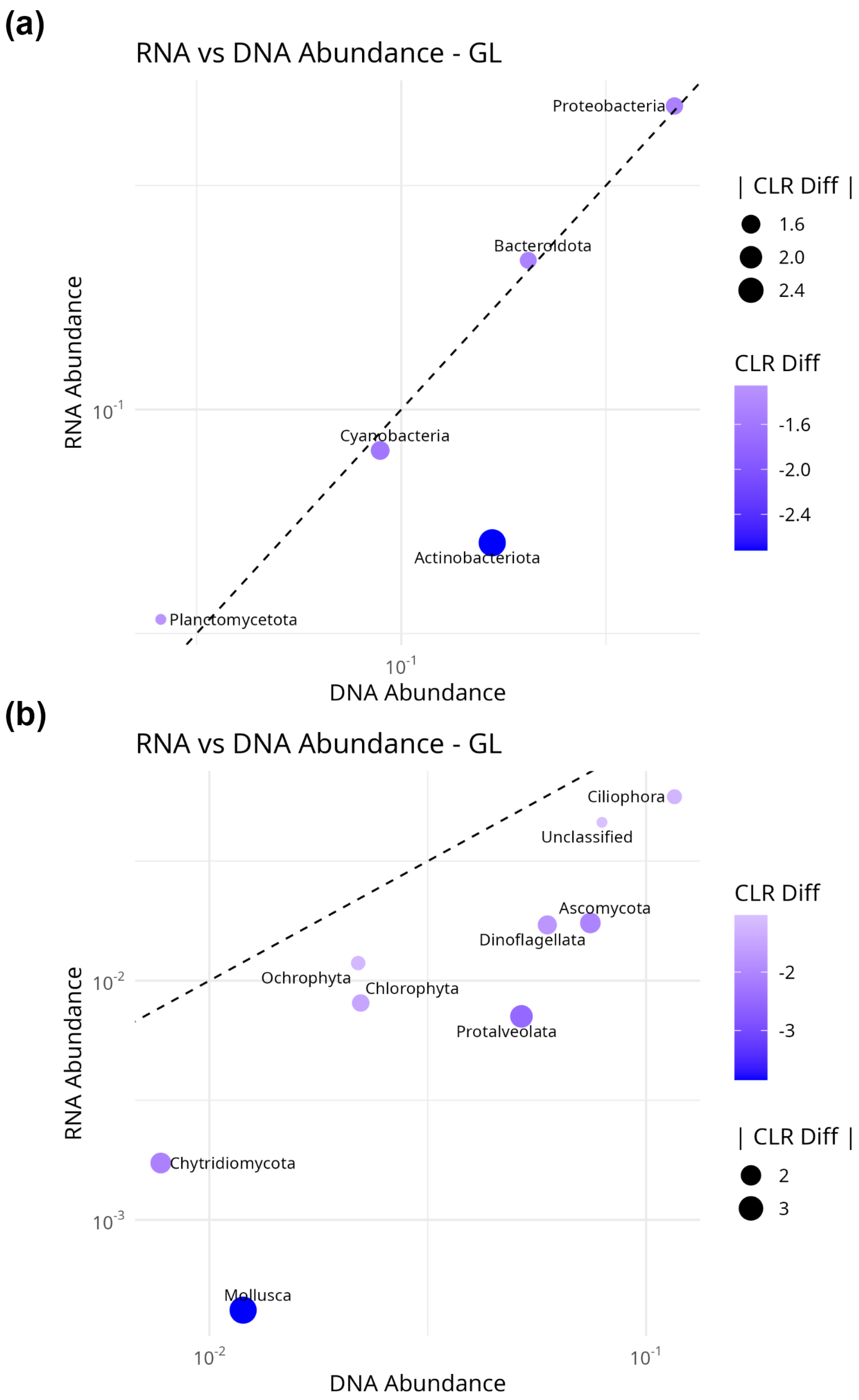


**FIGURE S10** Abundance and centered log-ratio comparisons for paired DNA and RNA samples for 16S families (a) and 18S phyla (b) from Grand Lake. x- and y-axes show the relative abundance of each taxon in the DNA and RNA samples respectively on a logarithmic scale. The size of each coloured dot indicates the absolute average difference between the CLR scores for paired RNA and DNA samples (see Methods), while the colour scale indicates the magnitude and direction of the difference. Red dots indicate CLR(RNA) > CLR(DNA), while blue dots indicate the opposite pattern. Only paired samples with absolute CLR difference greater than 1.0 are shown.

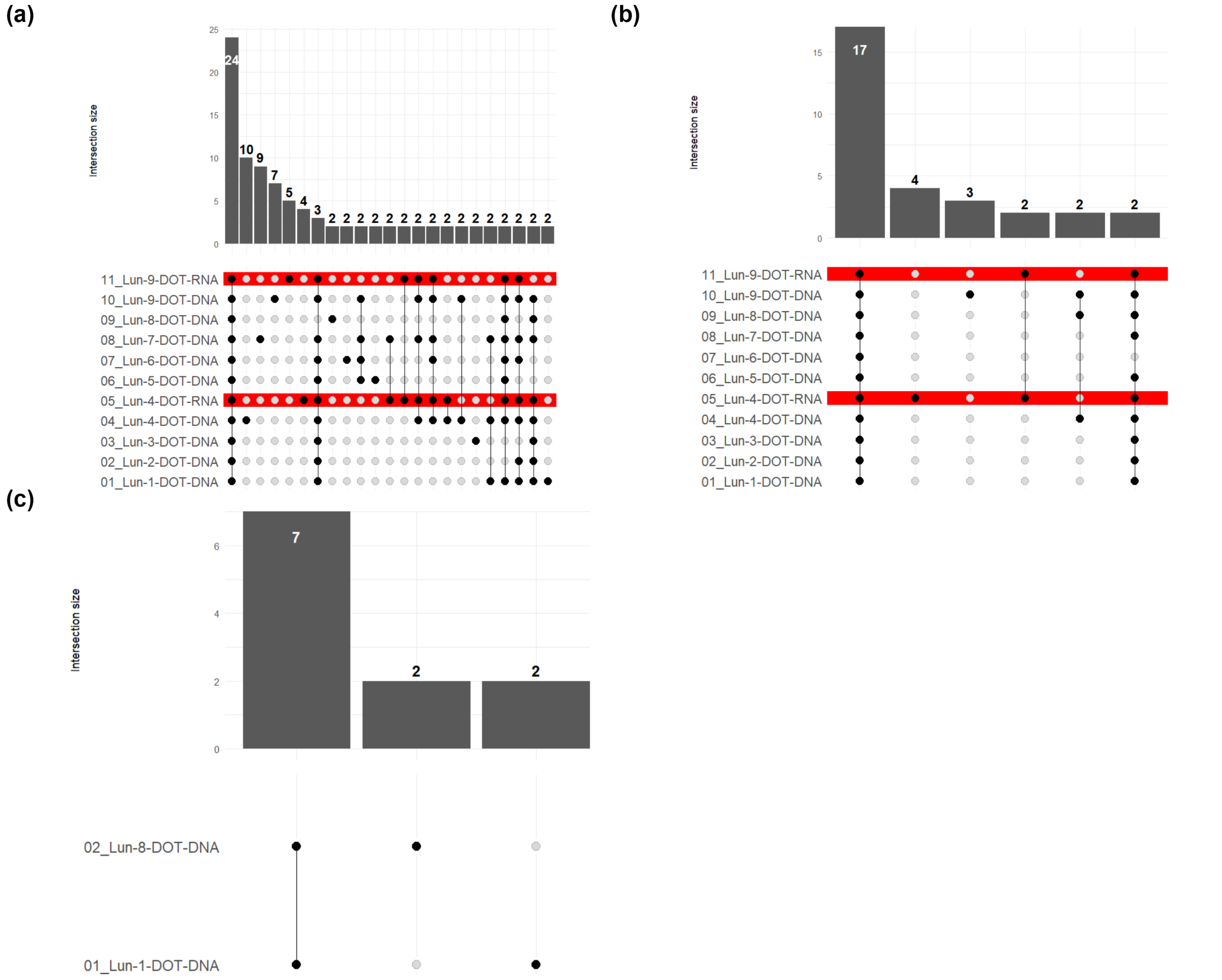


**FIGURE S11** UpSet plots showing distribution of taxa across all DOT DNA and RNA samples for 16S families (a), 18S phyla (b), and 12S species (c) profiles in Lunenburg Harbour. Each column in the plot indicates a unique presence / absence pattern across all samples, with black dots indicating presence and white dots absence. Bars at the top with numbers indicate the number of taxa that exhibit that presence/absence pattern. RNA profiles for 16S and 18S are highlighted in red. Numbers preceding each sample name are for ordering purposes only.

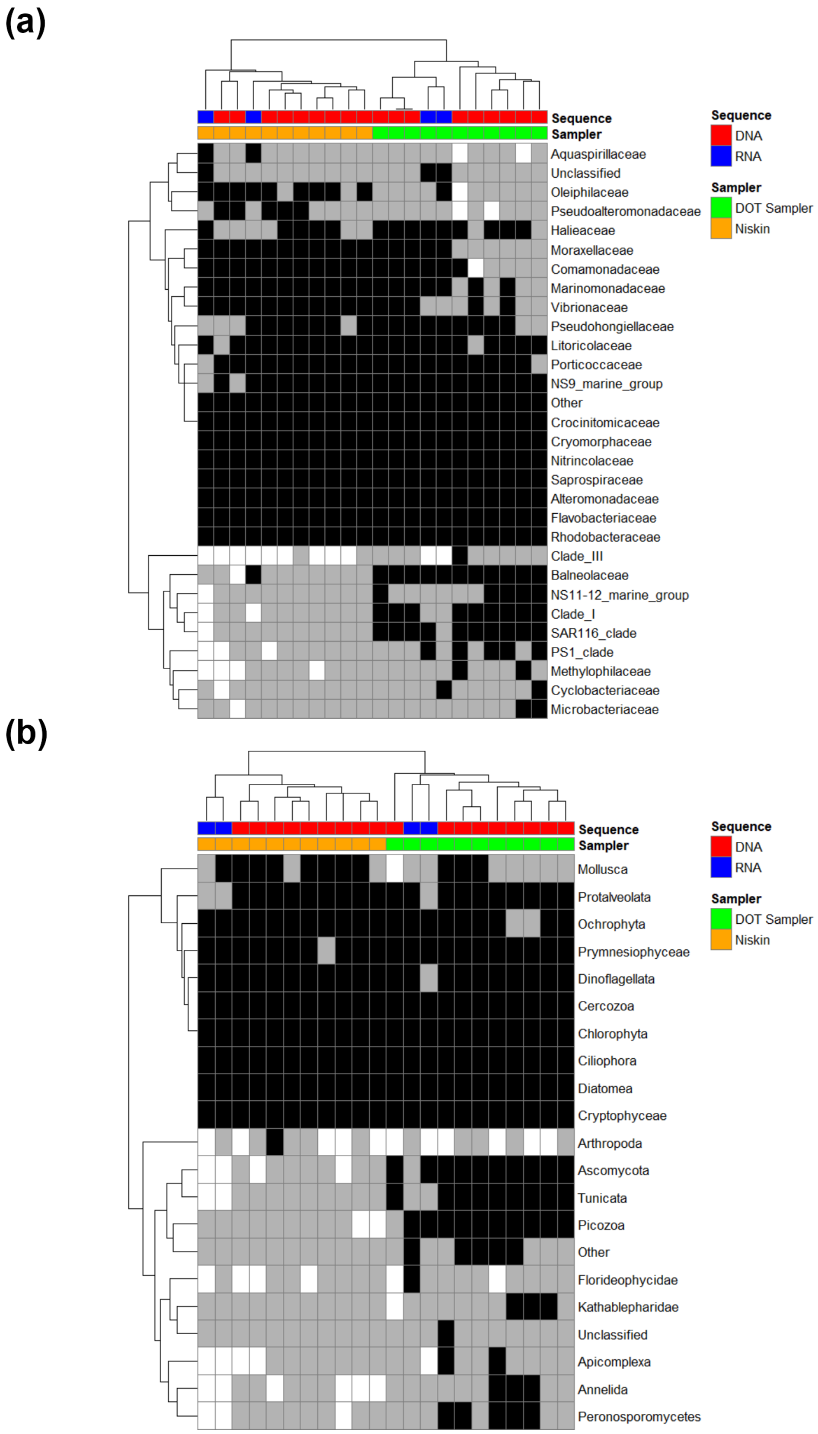


**FIGURE S12** Presence / absence heatmaps for all samples by sequence type and sampler for 16S families (a) and 18S phyla (b) from Lunenburg Harbour. All taxa shown are present in at least one sample at a minimum relative abundance of 1%. Samples with taxon abundance above this threshold are shown are indicated with black rectangles, while samples with taxon abundance less than 1% are shown in gray.

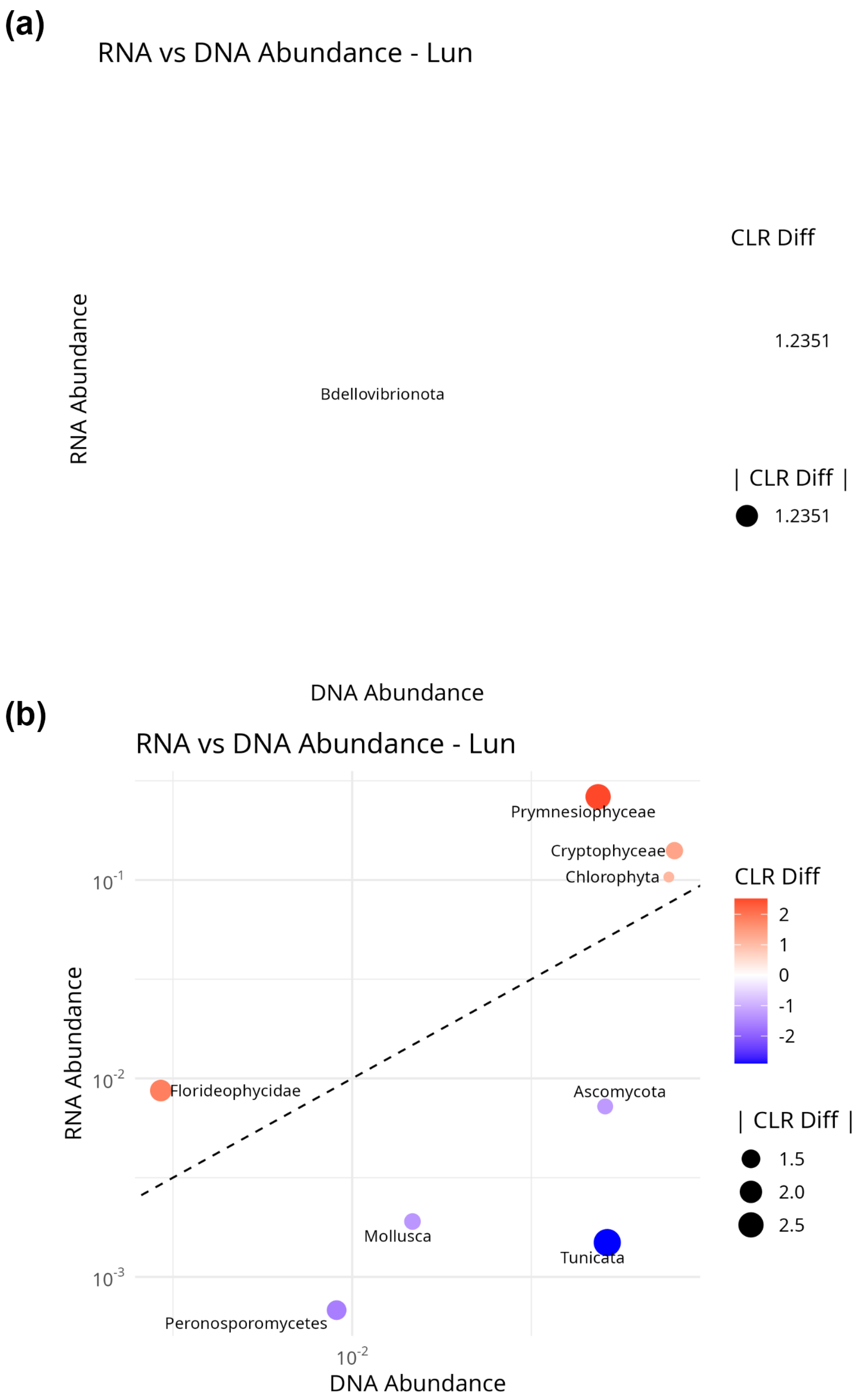


**FIGURE S13** Abundance and centered log-ratio comparisons for paired DNA and RNA samples for 16S families (a) and 18S phyla (b) from Lunenburg Harbour. x- and y-axes show the relative abundance of each taxon in the DNA and RNA samples respectively on a logarithmic scale. The size of each coloured dot indicates the absolute average difference between the CLR scores for paired RNA and DNA samples (see Methods), while the colour scale indicates the magnitude and direction of the difference. Red dots indicate CLR(RNA) > CLR(DNA), while blue dots indicate the opposite pattern. Only paired samples with absolute CLR difference greater than 1.0 are shown.

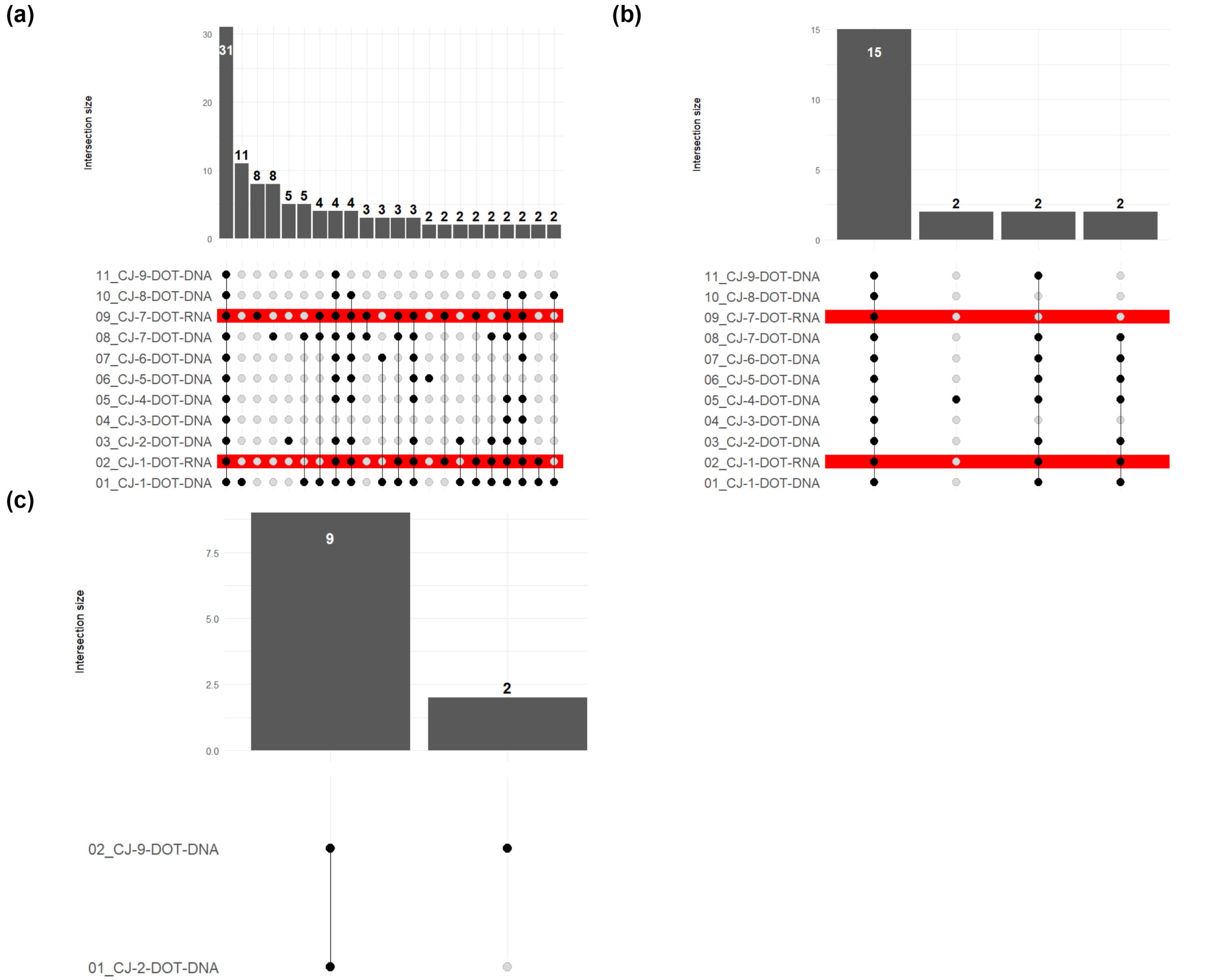


**FIGURE S14** UpSet plots showing distribution of taxa across all DOT DNA and RNA samples for 16S families (a), 18S phyla (b), and 12S species (c) profiles from the COVE Jetty in Halifax Harbour. Each column in the plot indicates a unique presence / absence pattern across all samples, with black dots indicating presence and white dots absence. Bars at the top with numbers indicate the number of taxa that exhibit that presence/absence pattern. RNA profiles for 16S and 18S are highlighted in red. Numbers preceding each sample name are for ordering purposes only.

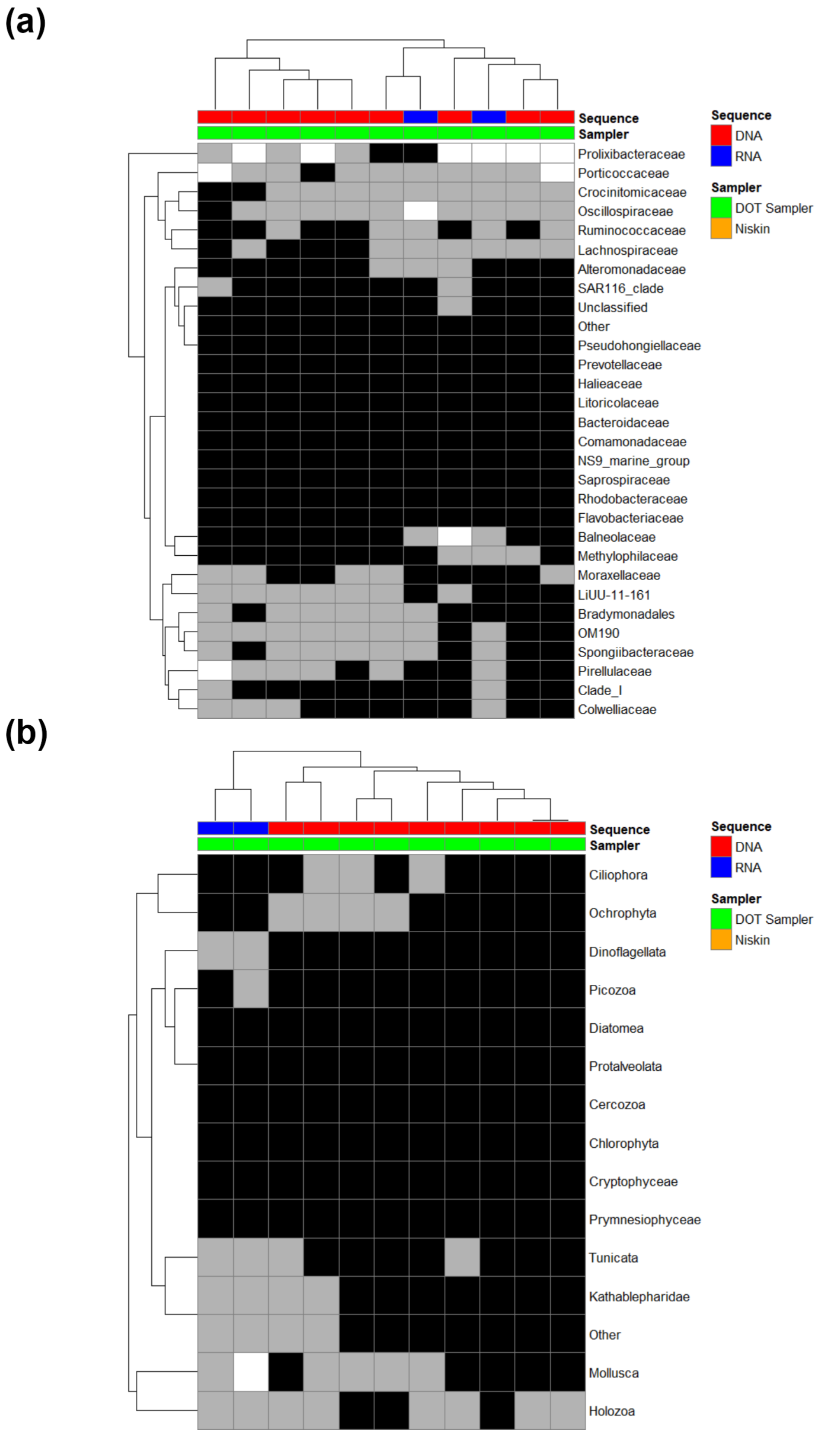


**FIGURE S15** Presence / absence heatmaps for all samples by sequence type and sampler for 16S families (a) and 18S phyla (b) from COVE Jetty. All taxa shown are present in at least one sample at a minimum relative abundance of 1%. Samples with taxon abundance above this threshold are shown are indicated with black rectangles, while samples with taxon abundance less than 1% are shown in gray.

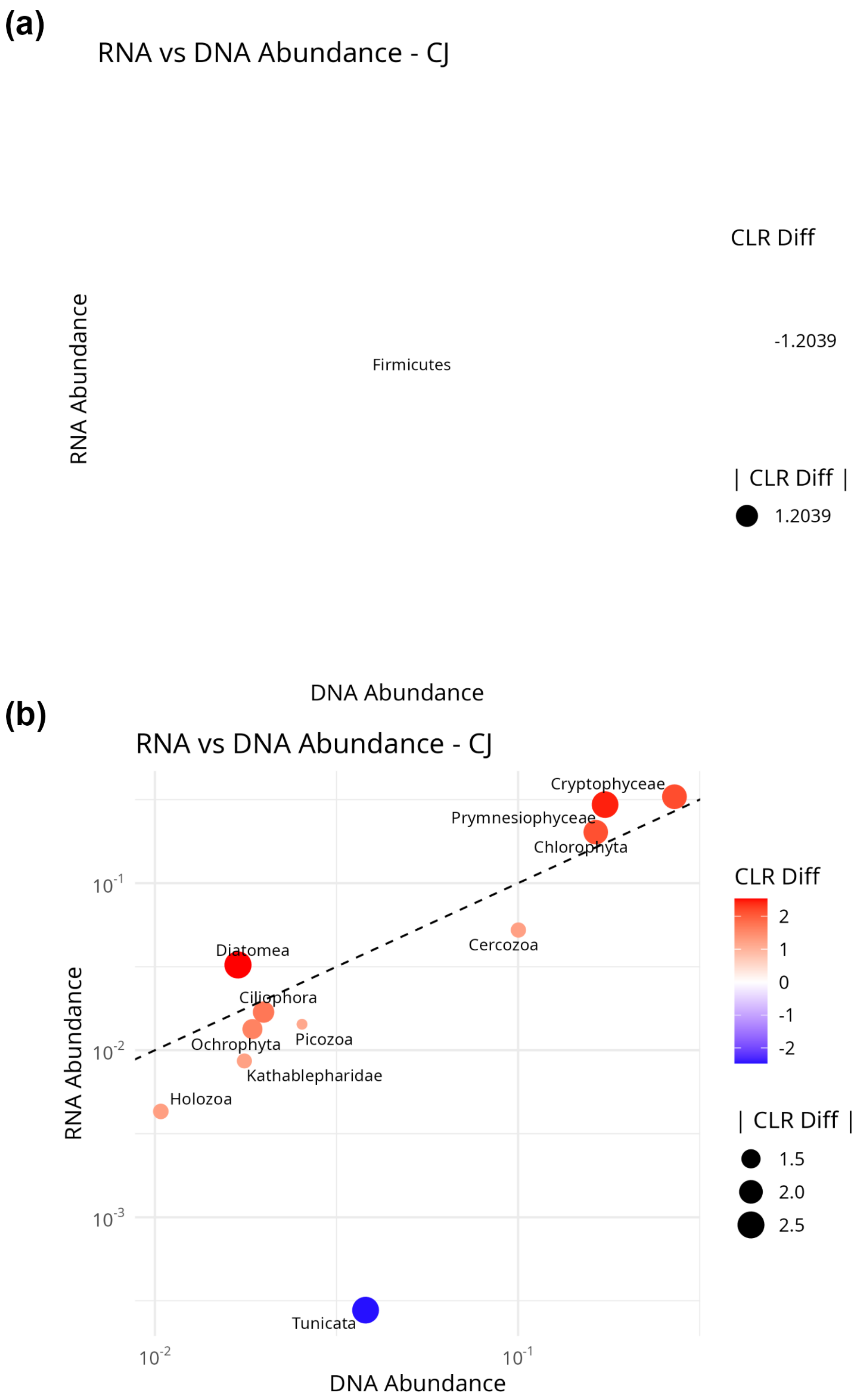


**FIGURE S16** Abundance and centered log-ratio comparisons for paired DNA and RNA samples for 16S families (a) and 18S phyla (b) from the COVE Jetty in Halifax Harbour. x- and y-axes show the relative abundance of each taxon in the DNA and RNA samples respectively on a logarithmic scale. The size of each coloured dot indicates the absolute average difference between the CLR scores for paired RNA and DNA samples (see Methods), while the colour scale indicates the magnitude and direction of the difference. Red dots indicate CLR(RNA) > CLR(DNA), while blue dots indicate the opposite pattern. Only paired samples with absolute CLR difference greater than 1.0 are shown.

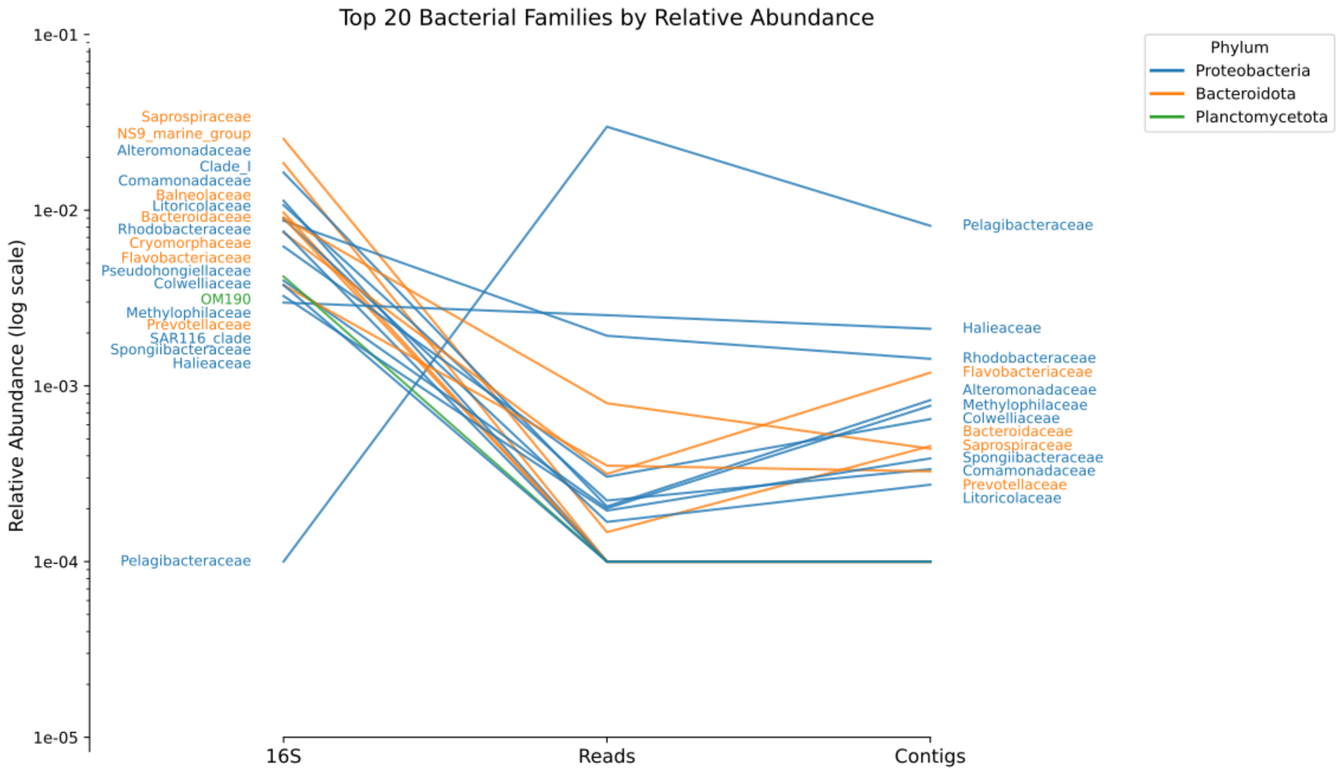


**FIGURE S17** Abundance profiles for the 20 most-abundant prokaryotic families inferred from 16S marker-gene and metagenomic reads and contigs from the COVE Jetty in Halifax Harbour. The y-axis indicates the relative abundance of a given family in each of the different sample types on a logarithmic scale. Colours indicate the phylum of each identified family.

**TABLE S1** Summary of samples, sequencing metrics, and metadata (Table S1).

| Location and Date | Sample Name | Method | Sample start time | Preservation time | Volume (mL) | [DNA] ng/µL | 16S DNA Reads | 16S cDNA Reads | 18S DNA Reads | 18S cDNA Reads | 12S DNA Reads | MG Reads |
|---|---|---|---|---|---|---|---|---|---|---|---|---|
| | SHP-1-DOT | DOT | 9:32 | 9:43 | 27 | 27.4 | 8698 | | 179 | | | 63387 |
| | SHP-2-DOT | DOT | 10:32 | 10:44 | 51 | 23.1 | 34313 | 93332 | 39910 | 118160 | | |
| | SHP-3-DOT | DOT | 11:32 | 11:44 | 33 | 18.0 | 13453 | | 12495 | | | |
| | SHP-4-DOT | DOT | 12:32 | 12:41 | 27 | 21.2 | 21598 | | 4873 | | | |
| Sheep's Head Pond | SHP-5-DOT | DOT | 13:32 | 13:45 | 45 | 19.9 | 19973 | | 3466 | | | 61237 |
| SHP | SHP-6-DOT | DOT | 14:32 | 14:45 | 39 | 15.8 | 17275 | | 9118 | | | |
| Tantallon, NS | SHP-7-DOT | DOT | 15:32 | 15:44 | 54 | 8.1 | 36305 | 103447 | 27390 | 119807 | | |
| 44°39'58.0"N | SHP-1-NIS | Niskin | 9:30 | 10:00 | 42 | 29.4 | 14255 | | 2138 | | | |
| 63°54'02.3"W | SHP-2-NIS | Niskin | 10:30 | 10:45 | 50 | 17.9 | 26773 | 84185 | 28095 | 113134 | | |
| July 17, 2023 | SHP-3-NIS | Niskin | 11:43 | 11:55 | 43 | 15.8 | 11718 | | 11381 | | | |
| | SHP-4-NIS | Niskin | 12:35 | 12:47 | 39 | 18.8 | 1 | | 0 | | | |
| | SHP-5-NIS | Niskin | 14:40 | 14:48 | 35 | 13.5 | 1 | | 9689 | | | |
| | SHP-6-NIS | Niskin | 13:43 | 13:50 | 42 | 13.2 | 6132 | | 8964 | | | |
| | SHP-7-NIS | Niskin | 15:34 | 15:43 | 43 | 5.3 | 32352 | 101011 | 32339 | 119403 | | |
| | GL-1-DOT | DOT | 9:47 | 9:57 | 30 | 10.1 | 4166 | | 5439 | | | 79331 |
| | GL-2-DOT | DOT | 10:32 | 10:38 | 39 | 12.1 | 5358 | | 2620 | | | |
| | GL-3-DOT | DOT | 11:17 | 11:23 | 42 | 5.8 | 2827 | | 6435 | | | |
| | GL-4-DOT | DOT | 12:02 | 12:08 | 39 | 12.1 | 6273 | | 5658 | | | |
| | GL-5-DOT | DOT | 12:47 | 12:54 | 45 | 4.0 | 10510 | 84112 | 7235 | 119537 | | |
| | GL-6-DOT | DOT | 13:32 | 13:39 | 48 | 9.4 | 188 | | 5482 | | | |
| Grand Lake | GL-7-DOT | DOT | 14:17 | 14:23 | 39 | 12.8 | 4532 | | 2911 | | | |
| GL | GL-8-DOT | DOT | 15:02 | 15:08 | 42 | 5.4 | 13027 | 74091 | 12483 | 107014 | | |
| Oakfield, NS | GL-9-DOT | DOT | 15:47 | 15:53 | 42 | 7.7 | 4603 | | 1495 | | 15814 | 82018 |
| 44°54'18.9"N | GL-1-NIS | Niskin | 9:50 | 10:03 | 52 | 4.8 | 4578 | | 322 | | | |
| 63°35'55.3"W | GL-2-NIS | Niskin | 10:33 | 10:40 | 63 | 8.4 | 56188 | | 26372 | | | |
| July 21, 2023 | GL-3-NIS | Niskin | 11:23 | 11:30 | 50 | 6.5 | 16369 | | 8778 | | | |
| | GL-4-NIS | Niskin | 12:07 | ~12:11 | 52 | 6.8 | 58019 | | 25225 | | | |
| | GL-5-NIS | Niskin | 13:13 | 13:17 | 43 | 3.4 | 11163 | 92666 | 5780 | 79018 | | |
| | GL-6-NIS | Niskin | 13:46 | 13:51 | 50 | 2.0 | 4270 | | 5456 | | | |
| | GL-7-NIS | Niskin | 14:30 | 14:38 | 55 | 2.9 | 6985 | | 5479 | | | |
| | GL-8-NIS | Niskin | 15:21 | 15:29 | 53 | 5.6 | 20085 | 95373 | 11833 | 95019 | | |
| | GL-9-NIS | Niskin | 15:57 | 16:02 | 50 | 5.2 | 4985 | | 10005 | | | |
| | Lun-1-DOT | DOT | 9:32 | 9:45 | 125 | 23.7 | 25389 | | 11293 | | | 70281 |
| | Lun-2-DOT | DOT | 10:17 | 10:28 | 125 | 13.6 | 21944 | | 12334 | | | |
| | Lun-3-DOT | DOT | 11:02 | 11:17 | 123 | 31.3 | 22824 | | 9696 | | | |
| | Lun-4-DOT | DOT | 11:47 | 12:02 | 120 | 14.6 | 41685 | 79337 | 33415 | 78848 | | |
| | Lun-5-DOT | DOT | 12:32 | 12:44 | 84 | 39.2 | 26884 | | 11790 | | | |
| | Lun-6-DOT | DOT | 13:17 | 13:27 | 69 | 42.1 | 23519 | | 5789 | | | |
| ABCO Dock | Lun-7-DOT | DOT | 14:02 | 14:15 | 96 | 29.0 | 33706 | | 4910 | | | |
| Lunenburg, NS | Lun-8-DOT | DOT | 14:47 | 14:57 | 72 | 31.4 | 17832 | 4927 | 30589 | 79258 | | |
| Lun | Lun-9-DOT | DOT | 15:32 | 15:43 | 78 | 13.2 | 37522 | 71379 | 35630 | 46355 | | |
| 44°22'31.4"N | Lun-1-NIS | Niskin | 9:39 | 9:58 | 30 | 111.9 | 34837 | | 8865 | | | |
| 64°19'01.3"W | Lun-2-NIS | Niskin | 10:18 | 10:26 | 40 | 39.7 | 25919 | | 12398 | | | |
| July 19, 2023 | Lun-3-NIS | Niskin | 11:12 | 11:18 | 20 | 36.2 | 32142 | | 37540 | | | |
| | Lun-4-NIS | Niskin | 11:57 | 12:06 | 19 | 8.0 | 35429 | 13581 | 37634 | 44050 | | |
| | Lun-5-NIS | Niskin | 12:35 | 12:40 | 28 | 38.0 | 28578 | | 22580 | | | |
| | Lun-6-NIS | Niskin | 13:33 | 13:39 | 26 | 30.0 | 16139 | | 23225 | | | |
| | Lun-7-NIS | Niskin | 14:20 | 14:25 | 30 | 29.0 | 35428 | | 17222 | | | |
| | Lun-8-NIS | Niskin | 14:57 | 15:03 | 28 | 24.4 | 21418 | | 21344 | | | |
| | Lun-9-NIS | Niskin | 15:41 | 15:49 | 31 | 4.4 | 21274 | 18502 | 34816 | 43868 | | |
| | CJ-1-DOT | DOT | 17:27 | 17:41 | 125 | 21.1 | 44065 | 75370 | 58755 | 121905 | | 14694509 |
| COVE Jetty | CJ-2-DOT | DOT | 20:02 | 20:15 | 96 | 39.6 | 14113 | | 41949 | | 80441 | 14968251 |
| CJ | CJ-3-DOT | DOT | 0:02 | 0:12 | 57 | 20.4 | 7688 | | 15147 | | | 15362084 |
| Dartmouth, NS | CJ-4-DOT | DOT | 4:02 | 4:19 | 125 | 24.1 | 23424 | | 43986 | | | 15863813 |
| 44°39'38.7"N | CJ-5-DOT | DOT | 8:02 | 8:12 | 125 | 15.6 | 6635 | | 35532 | | | 21703405 |
| 63°33'29.4"W | CJ-6-DOT | DOT | 12:02 | 12:11 | 125 | 12.1 | 18530 | | 32185 | | | 16692714 |
| July 22–24, 2023 | CJ-7-DOT | DOT | 16:02 | 16:13 | 84 | 16.2 | 50705 | 87261 | 62740 | 138934 | | 16655421 |
| | CJ-8-DOT | DOT | 20:02 | 20:16 | 111 | 45.4 | 18208 | | 20513 | | | 15297148 |
| | CJ-9-DOT | DOT | 0:02 | 0:15 | 125 | 44.0 | 9227 | | 29544 | | 66223 | 18890813 |

**TABLE S2** Statistics for MegaHIT assemblies (M1–M9). Genus names are based on Kraken2 annotations.

| MegaHIT Assemblies | M1 | M2 | M3 | M4 | M5 | M6 | M7 | M8 | M9 |
|---|---|---|---|---|---|---|---|---|---|
| # contigs (≥ 0 bp) | 6054 | 6657 | 6215 | 5549 | 6200 | 4336 | 4983 | 6149 | 7939 |
| # contigs (≥ 1000 bp) | 6054 | 6657 | 6215 | 5549 | 6200 | 4336 | 4983 | 6149 | 7939 |
| # contigs (≥ 5000 bp) | 1611 | 1567 | 1681 | 1261 | 1275 | 876 | 1210 | 1308 | 1385 |
| # contigs (≥ 10000 bp) | 492 | 562 | 615 | 418 | 422 | 329 | 478 | 524 | 460 |
| # contigs (≥ 25000 bp) | 104 | 134 | 156 | 108 | 156 | 120 | 143 | 164 | 121 |
| # contigs (≥ 50000 bp) | 43 | 43 | 60 | 36 | 45 | 37 | 55 | 50 | 44 |
| Total length (≥ 0 bp) | 31839659 | 35370067 | 37064402 | 28357715 | 31661455 | 22775388 | 29411453 | 32988181 | 36603097 |
| Total length (≥ 1000 bp) | 31839659 | 35370067 | 37064402 | 28357715 | 31661455 | 22775388 | 29411453 | 32988181 | 36603097 |
| Total length (≥ 5000 bp) | 18181451 | 20208360 | 23608131 | 15591115 | 17055547 | 12673295 | 18425537 | 18962901 | 17607163 |
| Total length (≥ 10000 bp) | 10625488 | 13419536 | 16304110 | 9860477 | 11395816 | 9059928 | 13419442 | 13663354 | 11480390 |
| Total length (≥ 25000 bp) | 5127757 | 7036005 | 9407010 | 5278395 | 7422923 | 5944302 | 8402881 | 8342435 | 6310551 |
| Total length (≥ 50000 bp) | 3009817 | 3940030 | 6193784 | 2869632 | 3660699 | 3040108 | 5348864 | 4339072 | 3756839 |
| # contigs | 6054 | 6657 | 6215 | 5549 | 6200 | 4336 | 4983 | 6149 | 7939 |
| Largest contig | 129130 | 194311 | 415927 | 156220 | 248731 | 177987 | 381104 | 200017 | 210960 |
| Largest contig Kraken2 | *Planktomarina* | *Planktomarina* | *Planktomarina* | *Pseudomonas* | *Flavobacterium* | *Flavobacterium* | *Alteromonas* | *Pseudomonas* | *Planktomarina* |
| Total length | 31839659 | 35370067 | 37064402 | 28357715 | 31661455 | 22775388 | 29411453 | 32988181 | 36603097 |
| GC (%) | 42.73 | 44.05 | 44.86 | 44.49 | 43.02 | 42.34 | 40.11 | 45.41 | 45.22 |
| N50 | 6010 | 6250 | 8016 | 5736 | 5623 | 5978 | 8350 | 6631 | 4749 |
| N90 | 2532 | 2470 | 2541 | 2448 | 2423 | 2423 | 2488 | 2428 | 2365 |
| auN | 15112.8 | 20687.5 | 32695.7 | 17322.3 | 20374.1 | 22154.2 | 32395.5 | 23034.3 | 17736.9 |
| L50 | 1197 | 1111 | 864 | 998 | 1044 | 641 | 620 | 878 | 1528 |
| L90 | 4632 | 5048 | 4527 | 4254 | 4756 | 3287 | 3636 | 4630 | 6237 |
| # N's per 100 kbp | 0 | 0 | 0 | 0 | 0 | 0 | 0 | 0 | 0 |

| Family | M1 | M2 | M3 | M4 | M5 | M6 | M7 | M8 | M9 | Total |
|---|---|---|---|---|---|---|---|---|---|---|
| Actinomycetales | | 2 | 1 | | | | | 1 | 1 | 5 |
| Alteromonadaceae | | 2 | 2 | | | | 2 | 2 | 3 | 11 |
| Enterobacteriaceae | | | 1 | | 2 | 1 | 1 | | | 5 |
| Flavobacteriaceae | 1 | 2 | 2 | | 2 | | 1 | 2 | 1 | 11 |
| Gammaproteobacteria | | | 1 | 1 | | 1 | 1 | | | 4 |
| Haliaceae | | | | | 1 | | | | | 1 |
| Litorivicinaceae | 1 | 1 | 1 | 1 | 1 | 1 | 1 | 1 | 1 | 9 |
| Micrococcales | | 1 | 1 | | | | 1 | 1 | | 4 |
| NA | | | | | 1 | 1 | | | | 2 |
| Pelagibacteriaceae | 3 | 1 | 3 | 1 | 1 | 1 | 3 | 1 | 3 | 17 |
| Rhodobacteraceae | | | 1 | | | 1 | 1 | | | 3 |
| Saprospiraceae | 2 | | 1 | 1 | 3 | 2 | | | | 9 |
| Vibrionaceae | | 1 | 1 | | | | 1 | 1 | | 4 |
| Total | 7 | 10 | 15 | 4 | 11 | 8 | 12 | 9 | 9 | 85 |

**TABLE S3** Distribution of families across samples M1–M9.

| Family | M1 | M2 | M3 | M4 | M5 | M6 | M7 | M8 | M9 |
|---|---|---|---|---|---|---|---|---|---|
| Actinomycetales | | 44.6 | 51.6 | | | | | 51.6 | 51.6 |
| Alteromonadaceae | | 61.0 | 34.5 | | | | 34.1 | 34.4 | 52.0 |
| Enterobacteriaceae | | | 100.0 | | 100.0 | 100.0 | 100.0 | | |
| Flavobacteriaceae | 36.1 | 33.4 | 33.2 | | 35.1 | | 31.3 | 33.0 | 38.5 |
| Gammaproteobacteria | | | 40.8 | 40.6 | | 40.6 | 40.6 | | |
| Haliaceae | | | | | 41.1 | | | | |
| Litorivicinaceae | 36.5 | 36.5 | 36.5 | 36.5 | 36.5 | 36.5 | 36.5 | 36.5 | 36.5 |
| Micrococcales | | 32.1 | 31.8 | | | | 32.6 | 31.8 | |
| NA | | | | | 33.8 | 33.8 | | | |
| Pelagibacteriaceae | 39.0 | 37.5 | 39.3 | 37.5 | 37.5 | 37.5 | 39.3 | 37.5 | 39.6 |
| Rhodobacteraceae | | | 33.9 | | | 33.9 | 33.9 | | |
| Saprospiraceae | 33.9 | | 31.2 | 31.2 | 36.6 | 39.4 | | | |
| Vibrionaceae | | 86.7 | 86.7 | | | | 86.7 | 86.7 | |

**TABLE S4** Relative abundances of families across samples M1–M9.

| Class | M1 | M2 | M3 | M4 | M5 | M6 | M7 | M8 | M9 | Total |
|---|---|---|---|---|---|---|---|---|---|---|
| disinfecting agents and antiseptics | 4 | 2 | 5 | 3 | 2 | 3 | 5 | 2 | 4 | 30 |
| elfamycin | | 1 | | | | | | 1 | 2 | 4 |
| fluoroquinolone; diaminopyrimidine; phenicol | | 1 | 1 | | | 1 | 1 | | | 4 |
| fluoroquinolone; tetracycline | | | | | | 1 | | | | 1 |
| glycopeptide | 3 | 5 | 7 | 1 | 5 | 3 | 5 | 5 | 3 | 37 |
| macrolide | | | | | 1 | 1 | | | | 2 |
| macrolide; streptogramin | | | 1 | | 1 | | 1 | | | 3 |
| rifamycin | | 1 | 1 | | | | 1 | 1 | | 4 |
| tetracycline | | | | | 1 | 1 | | | | 2 |
| Total | 7 | 10 | 15 | 4 | 11 | 8 | 12 | 9 | 9 | 85 |

**TABLE S5** Distribution of resistance classes across samples M1–M9.

| Family | M1 | M2 | M3 | M4 | M5 | M6 | M7 | M8 | M9 |
|---|---|---|---|---|---|---|---|---|---|
| disinfecting agents and antiseptics | 38.34 | 36.98 | 39.03 | 38.18 | 36.98 | 38.18 | 38.98 | 36.98 | 38.82 |
| elfamycin | | 87.02 | | | | | | 87.02 | |
| fluoroquinolone; diaminopyrimidine; phenicol | | 86.67 | 86.67 | | | 86.67 | 86.67 | | |
| fluoroquinolone; tetracycline | | | | | | 41.07 | | | |
| glycopeptide | 34.61 | 34.29 | 33.19 | 31.20 | 36.03 | 37.53 | 33.21 | 33.32 | 35.83 |
| macrolide | | | | | 100.00 | 100.00 | | | |
| macrolide; streptogramin | | | 100.00 | | 100.00 | | 100.00 | | |
| rifamycin | | 51.49 | 51.63 | | | | 51.56 | 51.56 | |
| tetracycline | | | | | 33.81 | 33.81 | | | |

**TABLE S6** Relative abundances of resistance classes across samples M1–M9.
